# Phylogenetic signal dynamics during niche filling in food webs

**DOI:** 10.1101/2025.11.04.686537

**Authors:** Alexandre Fuster-Calvo, Madelaine Proulx, Christine Parent, François Massol, Mathilde Besson, Paulo R. Guimarães, Dominique Gravel

## Abstract

Understanding how phylogenetic signal in ecological networks—the tendency for closely related species to resemble one another in ecological roles—emerges and persists remains a central challenge in community ecology. Here, we simulate food web evolution to track how the correspondence between phylogeny and trophic structure changes as communities assemble and niche space fills. By simulating trait evolution coupled with trait-matching for ecological interactions, we quantify how phylogenetic signal in trophic structure changes through time and examine how species’ network positions relate to their phylogenetic distinctiveness and diversification dynamics. We find that the signal declines over time, driven by emergent feedbacks between node extinction, link reorganization, and trait divergence. Species with high phylogenetic distinctiveness tend to be more specialized and occupy peripheral network positions, particularly in late-stage communities. Centrality consistently constrains diversification in intermediate consumers, emerges as a limiting factor for top predators after niche saturation, and shows nonlinear effects in basal species’ diversification. Applying our framework to empirical food webs from the Galápagos Islands, we find partial support for these predictions: phylogenetic signal in foraging and vulnerability roles declines with island age, but shows contrasting trends with island area and elevation. We also detect discrepancies between distance-based and clustering-based measures of phylogenetic signal, highlighting the need for robust methods to compare phylogenetic and network structures. Together, our results reveal how trophic interactions mediate the erosion of phylogenetic structure during community assembly and offer testable predictions for systems at different stages of diversification.

## Introduction

One of the most significant advances in community ecology has been the effort to link community structure with macroevolutionary processes (Webb et al. 2002, Mayfield & Levine 2010, Mouquet et al. 2012, Gerhold et al. 2015). Understanding how evolutionary history shapes ecological networks is increasingly recognized as essential for predicting network structure (Pearse & Altermatt 2013, Strydom et al. 2022) and anticipating how novel or perturbed communities will reorganize under global change (Rezende et al. 2007, Nuismer et al. 2018). In food webs, mutualistic, or host-parasite networks, a small set of phenotypic traits—such as body size or metabolic category—mediates interactions and explains much of the resulting network structure (Loeuille & Loreau 2009, Stouffer et al. 2011, Eklöf et al. 2013). These traits can be inherited from ancestral phenotypes and conserved across lineages, leading closely related species to share similar sets of interactions (Gómez et al. 2010).

Numerous studies have reported phylogenetic signal—i.e., phylogenetic correspondence without necessarily invoking a mechanistic process (Blomberg & Garland 2002)—in species interactions across diverse network types (Segar et al. 2020). Such a correspondence eventually scales up from pairwise interactions to entire network properties, e.g., modularity or inter-module connectivity (Peralta 2016), as well as species’ structural roles (Stouffer et al. 2012, Lai et al. 2021). However, many systems show no such correspondence (e.g., Ives & Godfray 2006, Rafferty & Ives 2013), and we still lack a general understanding of how phylogenetic signal in networks emerges and evolves (Brännström et al. 2012, Harmon et al. 2019, McGill et al. 2019).

Environmental filtering is expected to generate phylogenetic clustering when environmental tolerances are phylogenetically conserved—only lineages able to persist locally pass the filter (Mayfield & Levine 2010, Overcast et al. 2021). Beyond abiotic filtering, the role of biotic interactions in community assembly has been largely focused on competition (Cavender-Bares et al. 2009, Gerhold et al. 2015), with empirical evidence showing that it can generate density-dependent diversification (Rabosky 2013) and character displacement that reduces niche overlap (Schluter 2000). Classic niche-filling theory predicts that as communities assemble and niche space becomes saturated, competitive exclusion among close relatives should lead to phylogenetically overdispersed assemblages, thereby eroding phylogenetic signal (Darwin 1859, Gause 1934, Elton 1946, Diamond 1975, Webb 2000). However, biotic interactions beyond competition can also shape macroevolutionary trajectories by influencing speciation and extinction rates (Hui et al. 2018) and increasing resistance to species invasions (Tokita & Yasutomi 2003). For instance, classic hypotheses such as the Red Queen (Van Valen 1973) propose that predatory interactions drive adaptive trait evolution and diversification. Yet, these processes are typically studied in pairwise contexts (Hembry & Weber 2020), leaving open the question of how network-level structures affect diversification and community evolution. When accounting for multiple species and trophic levels, feedbacks and indirect effects emerge through competition and predation between them, shaping trait evolution in ways not predicted by pairwise interactions alone (Montoya et al. 2009, Schmitz 2010, Guimaraes et al. 2017).

To better understand how network and phylogenetic structures co-emerge, we need dynamic models that simulate trait evolution in tandem with species and interaction turnover, under the influence of different types of interactions (Caldarelli et al. 1998, Rossberg et al. 2005, Poisot & Stouffer 2016, Hembry & Weber 2020). Such models can generate explicit expectations for how phylogenetic signal arises and evolves during community assembly. Here, we adopt a simplified approach designed to isolate key mechanisms. Using a toy model of adaptive radiation and niche filling in which ecological opportunity drives rapid early diversification that slows as niche space becomes saturated (Stroud & Losos 2016), we simulate the evolutionary emergence of a food web and explore how phylogenetic signal changes over time.

We hypothesize that network phylogenetic signal will be strongest early in community assembly, when niche space is sparsely occupied and ecological opportunity (sensu predation/consumption) is high. Under these conditions, mutant lineages with similar trait values are more likely to establish successfully (Mayfield & Levine 2010), leading to clustered interactions among recently diverged species. As predicted by classic theory, this signal may weaken as niche space becomes saturated due to: (i) competitive exclusion among close relatives (Violle et al. 2011), and (ii) trait convergence among distantly related lineages occupying similar niches (Webb et al. 2002). In addition, we propose that network-mediated selection introduces novel drivers of phylogenetic signal evolution. Specifically, (iii) extinctions and its cascading effects through species interactions may disrupt the phylogenetic structure of the network in complex ways (Quince et al. 2005), and (iv) changes in network topology may feed back onto trait evolution by reshaping species’ ecological roles and the selective pressures they experience (Quince et al. 2002) (Figure 1ab). These emergent effects may generate unanticipated trajectories in the co-evolution of ecological and phylogenetic structure.

**Figure 1.**
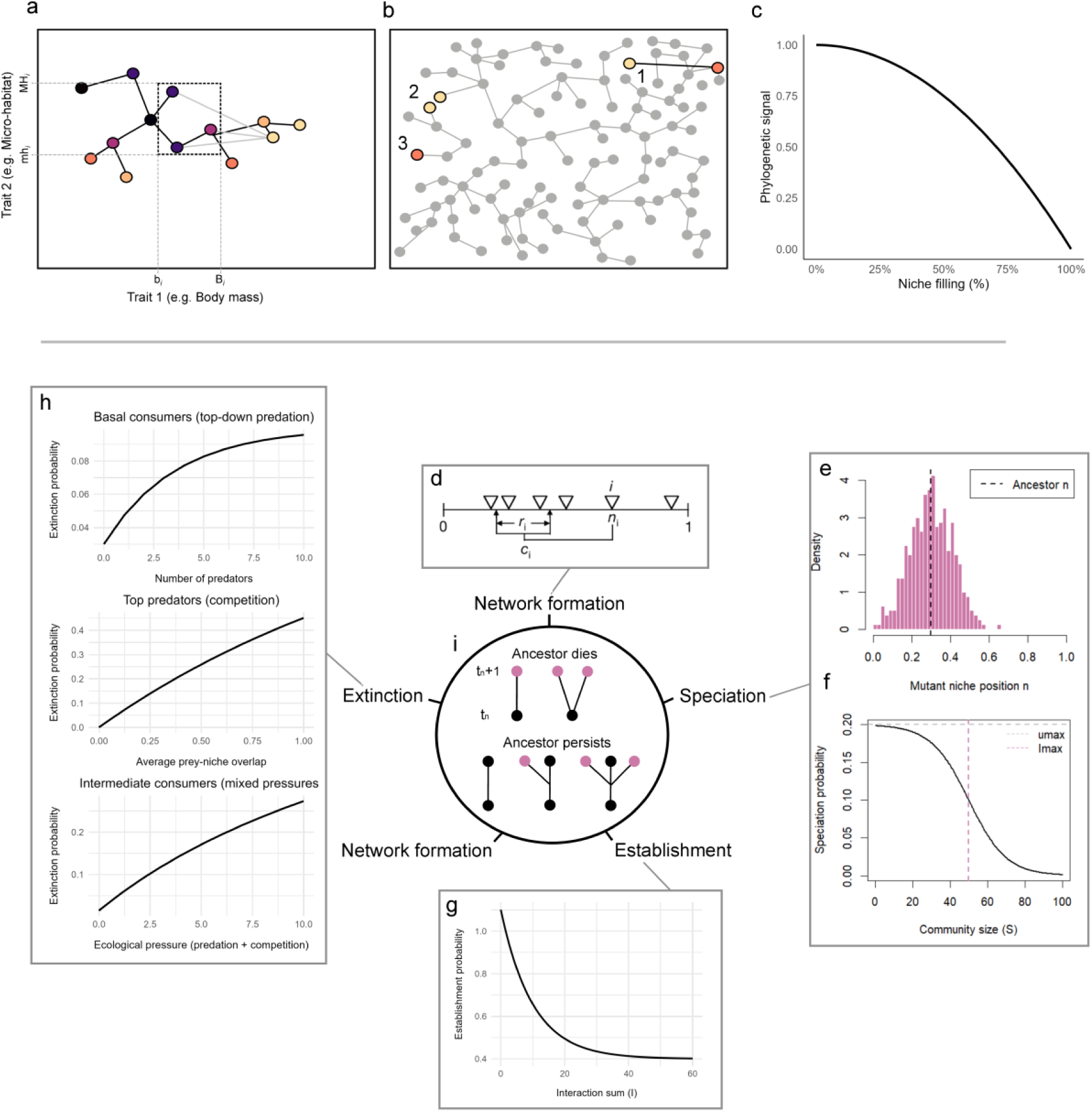
Conceptual hypothesis and model structure. (a-c) Hypothesis for the loss of phylogenetic signal as the community assembles and niche saturates. (a) Schematic niche space is delimited by two trait axes (rectangle). During early community assembly, when niche space is widely available (left panel), new species can establish via small trait shifts from ancestors, resulting in high phylogenetic signal. The possible diet range of species 𝑖’s diet is illustrated by the dashed rectangle. Light grey lines represent feeding links, and solid lines represent ancestral relationships. (b) In late-stage saturated niche space, phylogenetic signal is eroded due to: (1) successful establishment requiring large trait shifts to avoid competition, (2) ecological convergence of distantly related lineages, and (3) extinctions driven by network-mediated selection pressures, including predation, resource limitation, and indirect competition. (c) Prediction for the change in phylogenetic signal with increasing niche filling, as a consequence of processes in (a and b). Model structure (panels d–i): (d) Networks are built based on the niche model: species represented as triangles along the trait axis, each with an optimum and range, consuming a basal resource shown below the axis. (e) Speciation step illustrating the sampling of mutant trait values from the parental trait. (f) Speciation rate function declining with increasing community size, reflecting reduced diversification as niche space fills. (g) Establishment probability decreases in saturated communities. (h) Extinction probability as a function of ecological pressure depending on trophic role. (i) Schematic tree of ancestor-mutant branching outcomes, showing possible scenarios depending on ancestor survival and mutant establishment.

We further hypothesize that species’ positions in the phylogeny will help predict their ecological roles within the network. Specifically, species that are phylogenetically distinct from the rest of the community may possess more distinct ecological traits (Miranda & Parrini 2015, Dufour et al. 2024), enabling them to exploit prey types less used by other species and avoid competition to establish. As a result, they may tend to occupy more peripheral network positions, interacting with fewer species or engaging in specialized interactions. We predict that this negative association between phylogenetic distinctiveness and network centrality will intensify as niche space becomes saturated, reflecting stronger constraints on integration into the network core.

Finally, we expect that diversification dynamics will be shaped by species’ positions within the network, although the direction and strength of these effects may vary across trophic levels and community contexts. Basal consumers—herbivores that feed exclusively on basal resources—may experience increased diversification if predator richness promotes ecological differentiation, or a decrease if heightened predation imposes strong extinction (Vamosi 2005). Top predators—species with no consumers—may initially benefit from broad diets and central network positions, which buffer them and their descendants from extinction by reducing the risk of total prey loss (Gravel et al. 2011). However, as niche space saturates and competition among predators intensifies, these same traits may increase extinction risk and constrain diversification. Intermediate consumers—species that both consume and are consumed—are likely to be the most vulnerable, as they experience top-down control from predators and resource competition from other consumers. We therefore hypothesize that they will exhibit a hump-shaped diversification response, initially increasing with moderate prey richness but declining as predator richness and dietary overlap rise. This effect may weaken where predator richness generates positive indirect effects on intermediates.

Oceanic archipelagos offer ideal systems for exploring these questions (Warren et al. 2015). Their spatially discrete, relatively low-diversity communities serve as natural replicates along gradients of island age, area, and environmental complexity—factors that shape niche-filling dynamics during community assembly (Cardillo et al. 2008). In this study, our objectives are to (1) examine how the relationship between phylogenetic and network structure changes over time during food web evolution and niche saturation; (2) investigate how species’ positions in the network relate to their position in the phylogeny and their diversification dynamics; and (3) explore whether trends predicted by the model are qualitatively reflected in empirical data from vertebrate food webs of the Galápagos Islands.

## Methods

### Model functioning

We implemented a macroevolutionary assembly model in which lineages diversify by speciation and trait values shift at speciation in response to selection, with trophic interactions arising mechanistically from trait-matching rules (Figure 1c-h; see parameters’ summary in Table S1). This allows phylogenies and realistic network structures to emerge under explicit niche-filling dynamics. Following the niche model (Williams & Martinez 2000), each consumer species is defined by three continuous traits: its niche position (𝑛), its niche breadth (𝑟), and its foraging optimum (𝑐). Consumer interactions emerge mechanistically from trait matching: predators can consume prey whose niche positions fall within their foraging range defined by 𝑐 and 𝑟.

The food web structure emerges dynamically at each timestep from the traits of extant species. Each consumer’s foraging range is defined by its 𝑐 and 𝑟: [𝑐 − 𝑟, 𝑐 + 𝑟]. A consumer 𝑗 can consume a prey 𝑖 if the prey’s niche position 𝑛_𝑖_ lies within this range. Within its range, interaction probability with preys is determined by a Gaussian function of trait distance:

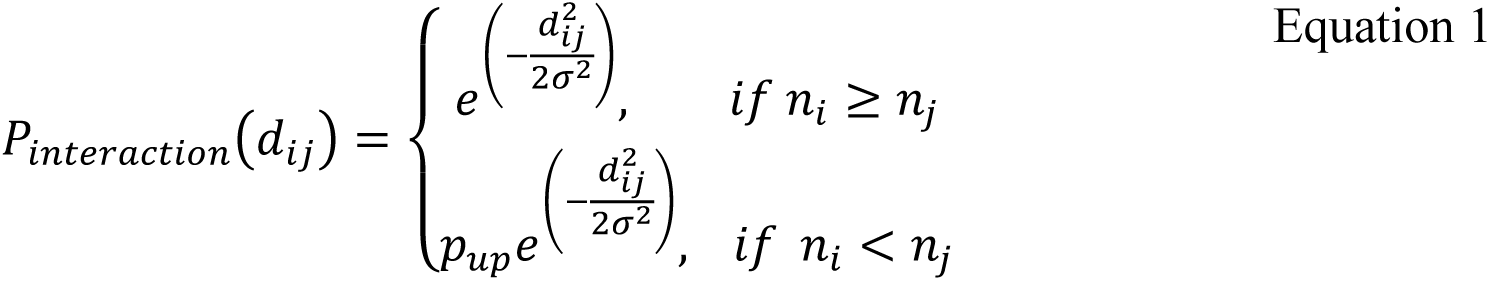

where 𝑑_𝑖𝑗_ is the distance between 𝑛_𝑖_ and 𝑐_𝑗_, and 𝜎 controls the interaction specificity. While 𝑟 defines the boundary beyond which interactions are not allowed, 𝜎 determines how interaction probability declines within this range. We keep 𝜎 fixed and independent of 𝑟 to allow species with similar foraging breadths to differ in diet selectivity. To impose a trophic hierarchy, we introduce a penalty factor 𝑝_𝑢𝑝_(set to 0.1) that reduces the interaction probability when a prey’s niche position exceeds that of its predator. Each potential interaction is realized stochastically and independently of all other interactions by drawing from a Bernoulli distribution with the computed probability. Self-feeding is explicitly disallowed by setting the diagonal of the interaction matrix to zero.

In the following, we always set the optimum 𝑐 at 𝑛⁄2, and we let both 𝑛 and 𝑟 evolve. The basal resource pool consists of 5 producers with niche positions drawn near zero from a uniform distribution on [0, 0.3], which do not evolve, to provide a minimal but structured resource base. This small variation allows early consumer lineages to diversify based on foraging traits, while ensuring a stable pool of basal resources throughout the simulation.

At each discrete timestep, each species has a chance to speciate, producing mutant descendants. The probability of speciation declines with increasing community size (following adaptive radiation theory; Yoder et al. 2010):

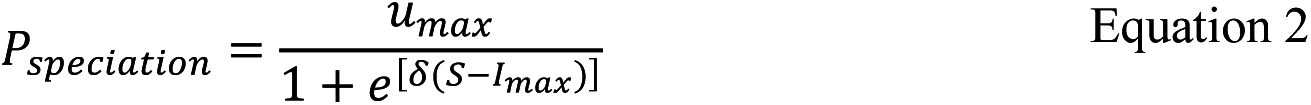

where 𝑢_𝑚𝑎𝑥_is the maximum speciation rate at low diversity, 𝛿 is a scaling parameter controlling the strength of density dependence, 𝑆 is the number of consumer (i.e., non-basal) species, and 𝐼_𝑚𝑎𝑥_represents the saturation point of the community. This threshold defines two phases in the simulation: the “before niche saturation” phase, in which speciation occurs more freely, and the “post niche saturation” phase, characterized by stronger diversity– dependent constraints on diversification. When speciation occurs, the ancestor produces two daughter species, each with modified traits. The first daughter species’ trait direction (greater or lesser) is chosen at random, and the second explores the opposite direction in trait space. Both 𝑛 and 𝑟 evolve through directional mutations generated from Beta distributions scaled around the ancestor’s value. The mutation process proceeds as follows: a direction (either greater or lesser) is selected, and the trait is displaced from the ancestor’s value by a scaled difference drawn from a Beta distribution whose shape depends on the trait value:

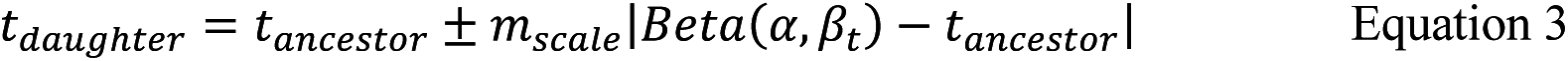

where 𝑡 represents either 𝑛 or 𝑟. The constant 𝑚_𝑠𝑐𝑎𝑙𝑒_is a scaling factor, set to 0.7, that moderates the magnitude of trait shifts during speciation. The shape of the Beta distribution depends on the ancestor’s trait value, with the first shape parameter defined as:

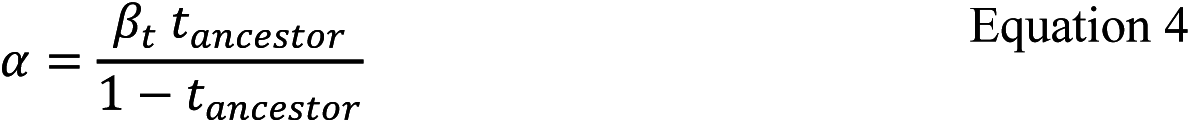

This ensures that the mean of the Beta distribution equals the ancestor

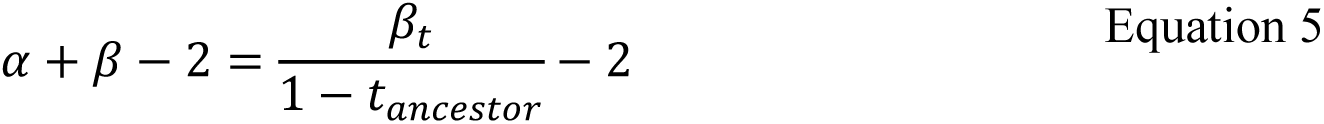

the distribution becomes more peaked around the ancestor’s value as the trait approaches the boundaries of its range (near 0 or 1). This narrows the range of possible mutations and constrains trait shifts when traits are extreme. Although trade-offs between niche position and niche breadth are often assumed (Williams & Martinez 2000, Loeuille & Loreau 2005), we allow these traits to evolve independently to avoid introducing additional assumptions and model complexity. Initial trait values are drawn independently from uniform distributions: 𝑛∼𝑢(0.1, 0.3) and 𝑟∼𝑢(0.05, 0.15).

After mutation, the establishment probability of the daughter species depends on the intensity of predation pressure from consumerscompetition and available niche space (Shea & Chesson 2002). To quantify this, we calculate the out-degree of the daughter species—i.e., the number of predators that would consume it, 𝐷_𝑜𝑢𝑡_. Establishment probability declines with increasing 𝐷_𝑜𝑢𝑡_, following:

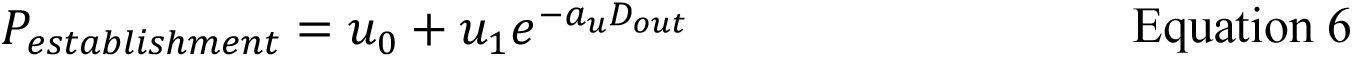

To prevent artificial persistence of functionally equivalent lineages, the ancestor goes extinct if the mutant’s niche position is too similar to its own, simulating exploitative competitive exclusion. This is implemented by comparing their niche positions and applying exclusion when the absolute difference is below a threshold parameter 𝜃_𝑛_, set to 0.05.

Survival of each species is determined probabilistically at each timestep, with extinction probabilities depending on the species’ position in the food web. Primary consumers— herbivores that feed exclusively on fixed basal resources—experience extinction risk driven by top-down predation pressure, modeled as a saturating function of the number of predators:

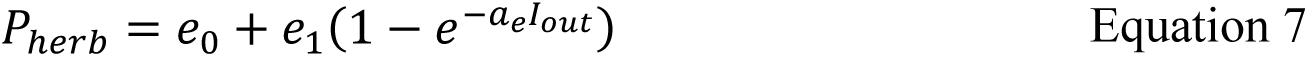

where 𝐼_𝑜𝑢𝑡_ is the number of predators consuming the herbivore. Top predators—species that are not consumed by any other—experience extinction risk that depends on bottom-up constraint and competitive similarity with other predators in the network. Predators that lose all their prey (i.e., have no outgoing trophic links) go extinct. The extinction risk also increases with overlap in prey use (dietary similarity), reflecting niche overlap and competitive exclusion dynamics:

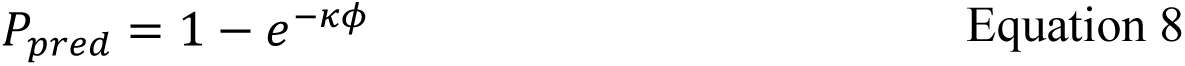

where 𝜅 is the competition coefficient controlling sensitivity to competitive overlap, and 𝜙 is the average trophic similarity of the species to others in the network. This is computed based on the network of trophic interactions among non-basal species 𝐿 at a given timestep (with entries 𝐿_𝑖𝑗_ = 1 if species 𝑖 consumes species 𝑗). We use a singular value decomposition (SVD) approach of this matrix (Dalla Riva & Stouffer 2016):

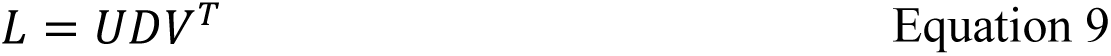

Note that in our model, species in the interaction matrix L act as prey in rows and as predators in columns, a setup we adopt for consistency with other components of the model (e.g., trait-based interaction matrices and extinction dynamics). This is the opposite convention to that used by Dalla Riva & Stouffer (2016), who define rows as predators and columns as prey. In this configuration, *V* contains species’ traits in predator-space (foraging profiles), *U* contains species traits in prey-space (vulnerability profiles), and *D* is the diagonal matrix of singular values (Rohr et al. 2010). Because we are interested in similarity in prey use (i.e., dietary overlap among predators), we focus on the rows of 𝑉. From this, we compute a similarity (or covariance) matrix between predators based on what they eat:

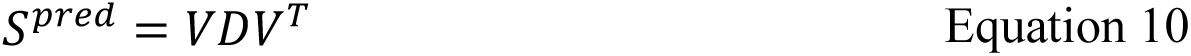

Here, 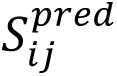 measures the degree of overlap in prey use between predator species 𝑖 and 𝑗. We then summarize average trophic similarity for each predator as the mean of its pairwise similarity scores:

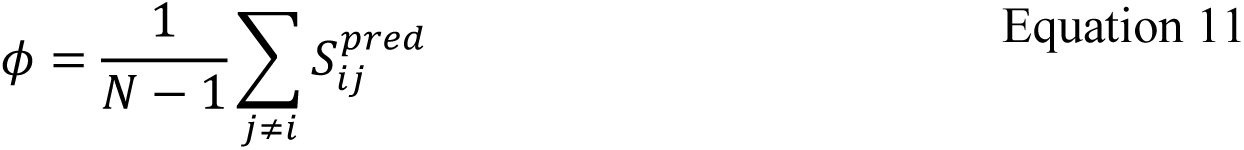

where *N* is the number of extant non-basal species. Intermediate consumers—species that both feed on other consumers and are themselves preyed upon—face extinction risk arising from both predation pressure and competitive similarity, modeled as a weighted combination:

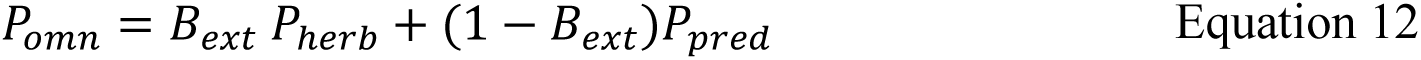

Where 𝐵_𝑒𝑥𝑡_ (between 0 and 1) balances the relative contribution of predation pressure versus competition. Species with no prey (i.e., top predators or intermediate consumers with zero outgoing links) are assigned an extinction probability of 1, reflecting obligatory dependence on prey resources.

At each timestep, we recorded species’ trait values, ancestor-descendant relationships (from which phylogenetic trees could be reconstructed), and interaction networks. We quantified temporal changes in network structure by computing connectance and modularity at each timestep. Connectance was calculated as the proportion of realized links out of all possible links, and modularity was computed using the Louvain algorithm (Blondel et al. 2008) on the undirected interaction matrix (as implemented in the *igraph* R package; Csárdi et al. 2025).

Simulations occasionally led to extinction cascades that caused the food web to collapse. We discarded these and retained only those that produced more than 30 species at the final timestep. A total of 344 simulations failed due to food web collapse, and 100 (23 %) successful runs were retained for analysis.

### Relating network and phylogenetic structures

We evaluated the correspondence between network and phylogeny structures. We represented species as multivariate trait vectors in both phylogenetic and trophic network spaces. For the phylogeny, we first computed the variance-covariance matrix of each tree— representing expected trait covariation under Brownian motion (Garland & Ives 2000)— using the *vcv* function in the *ape* R package (Paradis & Schliep 2019). We then scaled this matrix to a correlation matrix and performed an eigen-decomposition to obtain phylogenetic trait scores. For the trophic network, we performed singular value decomposition (SVD) of the trophic interaction matrix, yielding trait representations of species based on their vulnerability or prey (𝑈√𝐷) and foraging or predator (𝑉√𝐷) profiles (Rohr et al. 2010, Dalla Riva & Stouffer 2016; note that this follows the opposite convention, where species are represented as rows = prey and columns = predators).

#### Mantel correlation assessment

We assessed the correspondence between phylogenetic and network-derived species positions with Mantel correlations at each timestep (Mantel 1967, Cattin et al. 2004, Fontaine & Thébault 2015). For each role (predator and prey), we used the species scores from the trophic decomposition and from the phylogenetic eigen-decomposition. We retained enough leading axes to explain 80% of the cumulative variance in both network and phylogenetic spaces, ensuring comparable dimensionality. We then computed pairwise Euclidean distances among species in each space. Finally, we applied a Mantel test (with 999 permutations) between the phylogenetic distance matrix and the corresponding network-derived distance matrix using the *vegan* R package (Oksanen et al. 2013), obtaining a correlation coefficient quantifying phylogenetic signal in trophic structure.

#### Clustering similarity assessment

We compared species clustering inferred independently from the phylogeny and from the trophic interaction network at each timestep. We converted the variance-covariance matrix of the phylogenetic tree into an unnormalized graph Laplacian (Chung 1997), a matrix that emphasizes contrasts in trait covariance among species. This transformation allows the use of spectral clustering (Ng et al. 2001), a method that partitions species into groups based on shared structure in the Laplacian’s eigenvectors—in this case, species with similar phylogenetic positions in trait covariance space.

We determined the number of phylogenetic clusters with a spectral gap heuristic, which identifies the largest significant jump between successive eigenvalues of the Laplacian. The idea is that the optimal number of clusters corresponds to the point beyond which additional dimensions (i.e., clusters) offer diminishing returns in explanatory power. Once this number was identified, we applied spectral clustering to assign each species to a phylogenetic cluster.

We applied a Bernoulli stochastic block model (SBM) to estimate both the number of groups and the membership probabilities, yielding discrete cluster assignments for each species within the interaction network (food web). SBM estimation was performed using the *sbm* R package (Chiquet et al. 2021).

We quantified the agreement between phylogenetic clustering and interaction network-based clustering with the Normalized Mutual Information (NMI) (Danon et al. 2005) between the two cluster assignments at each timestep, using the *compare* function of the *igraph* R package. NMI measures shared information between partitions, ranging from 0 to 1, with higher values indicating stronger alignment of phylogenetic and network-derived species groups (Astegiano et al. 2017).

### Testing for size-driven artifacts via randomization

Because the number of species in the food webs and phylogenies changes over time in our simulations, the observed trend of correlations with time could be driven by variation in network or tree size, rather than ecological or evolutionary signal. To control for this, we implemented a randomization procedure for each metric at each timestep (Mouquet et al. 2012, Perez-Lamarque et al. 2022). Specifically, we performed 100 randomizations per timestep by permuting species identities in the network-derived distance matrices for the Mantel tests, and in the cluster assignments for the NMI measures, while keeping the phylogenetic distance matrix fixed. This preserved the structure of the phylogeny while removing any correspondence between phylogenetic relatedness and interaction similarity. Observed Mantel correlations and clustering NMI values were then visually compared against the resulting null distributions to assess whether phylogenetic signal trend with time deviated from random expectations.

### Relationship between species’ phylogenetic distinctiveness and interaction degree

We explored whether phylogenetic distinctiveness (PD) is associated with species’ trophic roles in the network. At each timestep, we computed species’ PD as the fair proportion index using the recorded phylogenetic tree with positive branch lengths (function *evol.distinct* from the *picante* R package; Kembel 2010). Consumer niche breadth was measured as each species’ in-degree (Freeman 1977) in the directed interaction network, representing the number of prey types it consumes. We excluded basal consumers from the analysis to focus on active consumer roles. These values were then combined per timestep to evaluate the relationship between phylogenetic distinctiveness and consumer niche breadth.

### Relationship between diversification rates and network centrality

We explored how network position relates to diversification dynamics by tracking speciation, extinction, and network centrality metrics for each species across simulations. For species in each replicate, we recorded their origination time, extinction time, and number of descendant species. From these records, we calculated species-level diversification metrics summarized over their entire evolutionary lifespan: speciation rate (descendants per unit time), extinction rate (inverse of longevity), and net diversification (speciation minus extinction).

To examine how network position relates to diversification within these trophic roles, we selected metrics that reflect different ecological pressures relevant to each trophic guild. For basal consumers, we used out-degree (number of predators; Freeman 1977) to capture direct predation risk, and eigenvector centrality to account for their importance by considering both direct links and the centrality of their predators (Bonacich 1987)—to explore the consequences of being consumed by more generalist or specialist predators. For intermediate consumers and top predators, we used in-degree as a measure of resource breadth, and betweenness centrality to reflect their role as connectors linking different parts of the food web—i.e., species that lie on many shortest paths and may regulate energy flow between otherwise separate modules (Freeman 1977). We computed these centrality metrics and trophic level for all species at each timestep, then summarized them by taking the mean over their evolutionary lifespan. We standardized (z-scored) all centrality measures within each timestep of each simulation to compare across simulations with various network sizes and structures. All centrality calculations were performed using the *igraph* package.

### Galápagos food web data

We compared model predictions with a food web from the Galápagos Islands. This system has clearly defined geographic boundaries, a shared regional species pool, and strong environmental gradients likely to influence niche availability. We used a dataset compiled by Proulx (2022) on vertebrate predator-prey interactions across the archipelago. This data consists of a regional metaweb spanning 13 islands, with a mean age of 0.8 million years [range: 0.04, 3 mya], mean area of 145 km² [range: 15 to 4726 km²], and mean elevation of 397 meters [range: 67 to 1685 m]. The interaction data was compiled through an extensive literature review, incorporating published reports, articles, and expert knowledge. Species lists for each island were assembled primarily from the Charles Darwin Foundation database (https://datazone.darwinfoundation.org/en/checklist) and GBIF records, with systematic correction of taxonomic inconsistencies. The diet of all terrestrial vertebrates was compiled with an extensive literature search, including scientific articles, reports, and other resources accessible on the web. All pairwise interactions were collated in a metaweb, and local (island) webs were extracted by subsetting co-occurring species only. Proulx (2022) also assembled a phylogenetic tree for Galápagos vertebrate predators by combining topologies from the TimeTree database (Kumar et al. 2017) with taxon-specific phylogenies (Figure S1).

We applied the same methods used in the simulations to test for phylogenetic signal in trophic networks and assess the relationship between phylogenetic distinctiveness and species’ trophic positions. To explore whether trends consistent with our model predictions are present, we qualitatively examined how these relationships vary with island characteristics likely to influence niche saturation. Specifically, we considered three metrics approximating varying niche saturation states: island age (older islands may support greater diversity and functional assembly), island area (larger areas offer more niche availability), and island elevation (topographic complexity increases niche diversity). Island ages were obtained from Geist et al. (2014) and area and elevation from Weigelt et al. (2013).

## Results

Extinction risk, trophic structure, and network organization change significantly over the course of community assembly. Competition-driven extinction probability was highest early in the simulations and declined over time (Figure 2a). Top-down extinction probability peaked early for basal consumers and then declined, while it progressively increased for intermediate consumers until stabilizing after the community reached its carrying capacity (Figure 2b). In contrast, the number of bottom-up extinctions increased throughout the simulations, while the mean per-species extinction probability decreased and then stabilized (Figure 2c). Range trait values varied across trophic levels, increasing over time for top predators and basal consumers, but decreasing after saturation for intermediate consumers (Figure 2d). The number of species per trophic level increased steadily until community size saturated: top predators reached a plateau, intermediate consumers declined, and basal consumers continued to increase (Figure 2e). Food web connectance declined over time, while modularity increased (Figure 2f–g).

**Figure 2.**
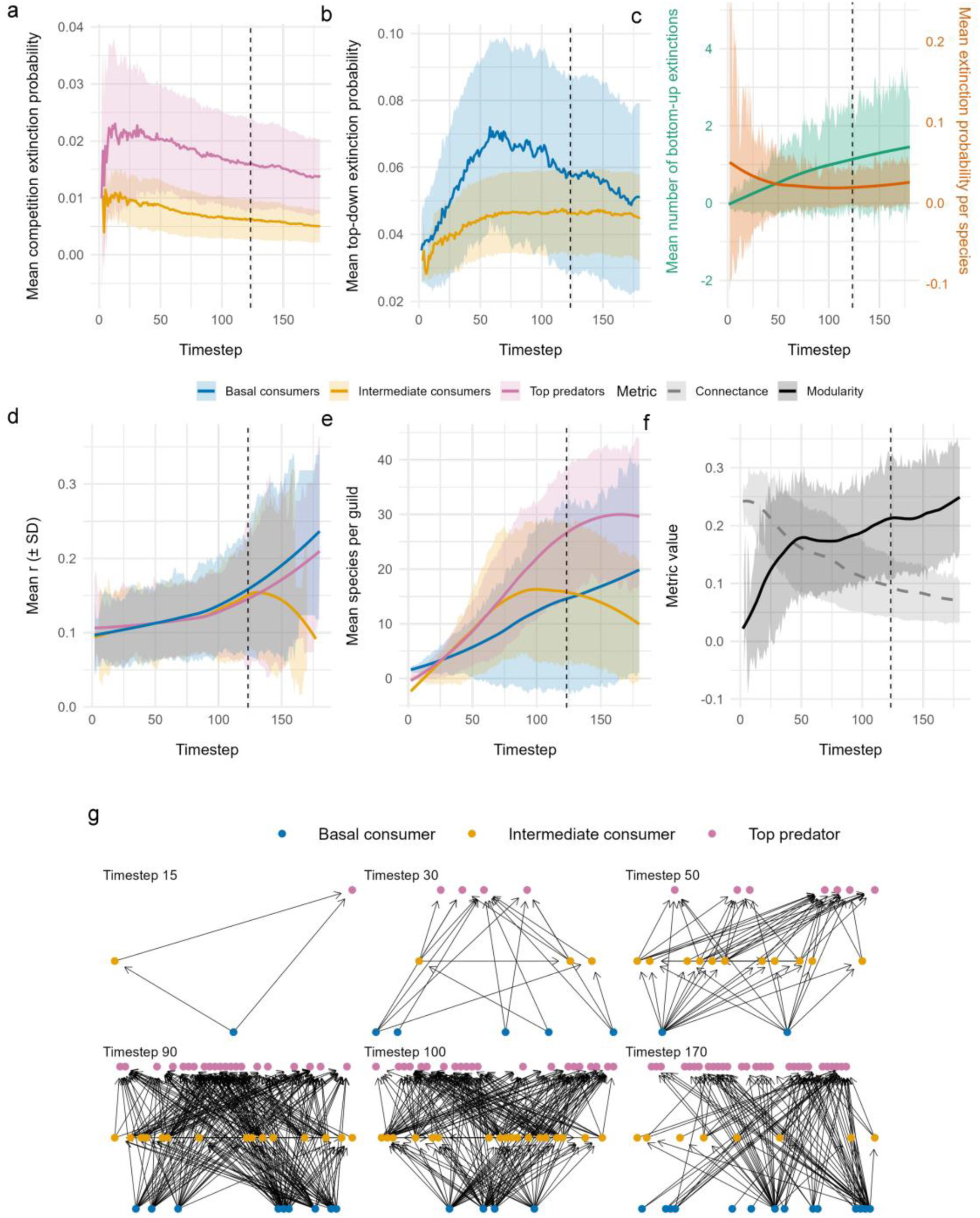
Dynamics of extinction probability, trophic structure, and network organization during simulations. Species turnover in the simulations is driven by speciation and extinction dynamics, with community size increasing over time until reaching the carrying capacity 𝑆_𝑚𝑎𝑥_ = 60 (indicated by the dashed vertical line). Panels (a–c) show the mean contribution of different processes to extinction events over time. (a) Mean competition-driven extinction probability. (b) Mean top-down extinction probability, driven by predator pressure. (c) Mean number of bottom-up extinctions and per-species extinction probability. (d) Mean range trait across trophic levels. (e) Mean number of species per trophic level. (f) Network connectance and modularity over time. Panel (g) illustrates the evolution of network architecture for one example simulation.

Phylogenetic signal declined over time with the assembly of the community. Mantel correlation between phylogenetic distances and network trait distances showed a rapid decline during the early stages of community assembly, then declined more gradually, eventually approaching a plateau once niche space was saturated. Normalized mutual information (NMI) between phylogenetic and network-based clusters decreased progressively over time (Figure 3).

**Figure 3.**
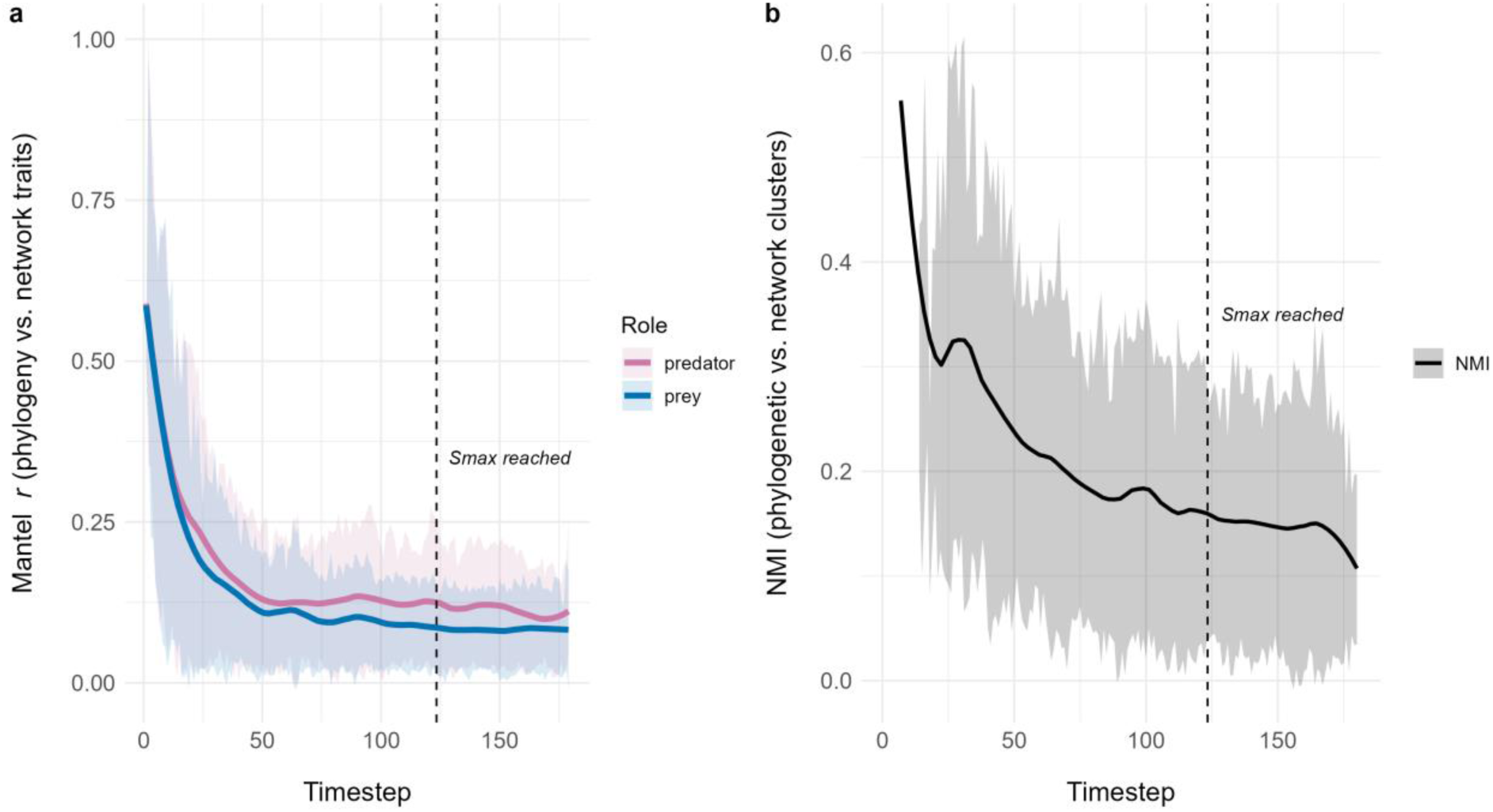
Phylogenetic signal during niche filling. We quantified phylogenetic signal over time using two approaches. (a) Mantel correlation between pairwise distances among species in phylogenetic trait space (from eigen- decomposition of a phylogenetic correlation matrix) and in trophic trait space (from singular value decomposition of predator foraging or prey vulnerability profiles). (b) Normalized Mutual Information (NMI) between clusters inferred independently from the phylogeny (via spectral clustering of a trait covariance Laplacian) and from the interaction network (via SBM clustering). Solid lines show LOESS-smoothed means, and shaded areas represent ±1 SD across simulations.

Phylogenetic distinctiveness (PD) was negatively related to species in-degree (i.e., diet breadth), both before and after niche saturation. In both phases, species with low to intermediate PD exhibited a wide range of in-degree values, including the most generalist species. However, as PD increased, the spread of in-degree narrowed, converging toward low values at high PD (Figure 4).

In top predators, diversification declined nearly linearly with increasing in-degree (diet breadth) before niche saturation, but after saturation, the relationship became hump-shaped, with the highest diversification at intermediate diet breadth. Intermediate consumers showed a similar relationship. In contrast, betweenness centrality showed no clear relationship with diversification in top predators before saturation. After saturation, diversification peaked at low betweenness, but became highly variable and tended to decline at higher values. For intermediate consumers, diversification declined linearly with increasing betweenness in both phases (Figure 5a-d). For basal consumers, diversification declined almost linearly with increasing out-degree (number of consumers) in both phases. For eigenvector centrality, diversification followed a hump-shaped relationship, with rates peaking at intermediate to high centrality, and remaining relatively elevated at the upper end (Figure 5e-h).

**Figure 4.**
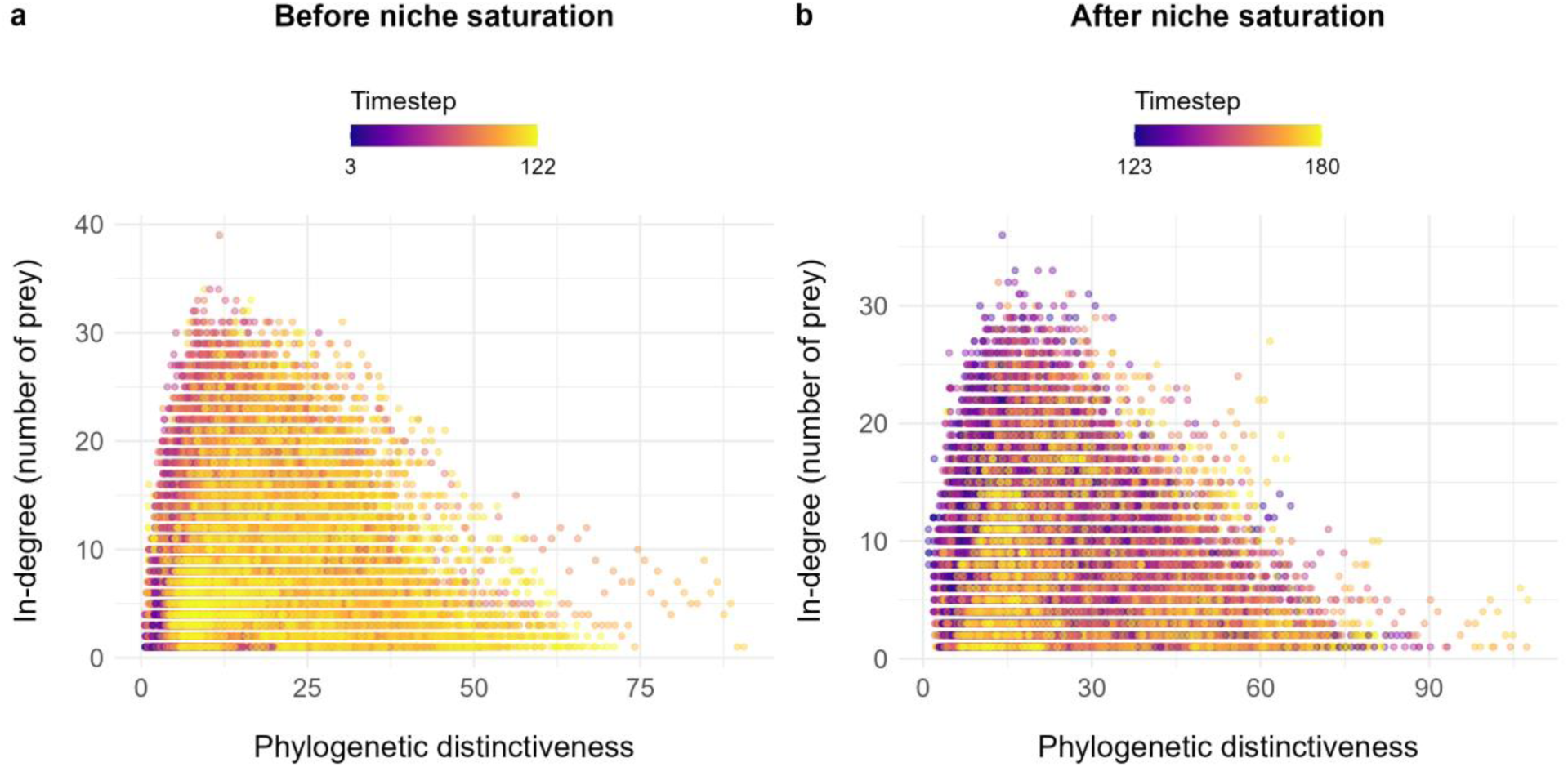
Phylogenetic distinctiveness and species’ diet breadth. Relationship between phylogenetic distinctiveness and species’ in-degree (i.e., diet breadth) across all timesteps in 100 simulations. (a) Before the niche saturation phase, i.e., prior to the community reaching its carrying capacity of 60 species, and (b) after the saturation phase.

**Figure 5.**
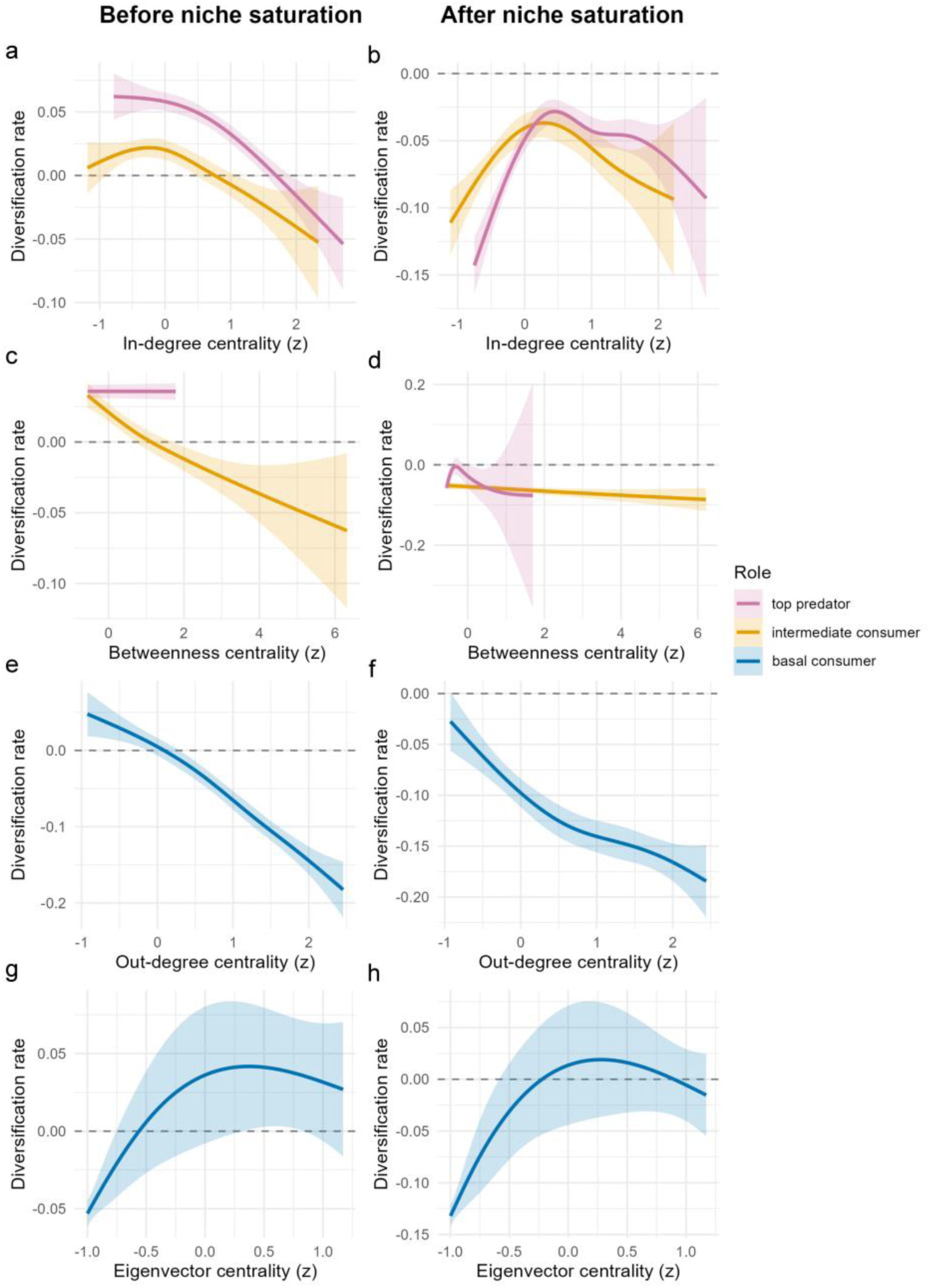
Relationship between species’ diversification rates and network centrality. The left column shows trends before niche saturation (i.e., community carrying capacity of 60 species), while the right column shows relationships after saturation. Top predators (species feeding on others but not consumed themselves) and intermediate consumers (species that both feed on others and are consumed) are shown in panels (a-d). Panels (a, b) display out-degree, defined as the number of prey species a consumer feeds on, and panels (c, d) show betweenness centrality, which captures how often a species lies on the shortest paths connecting others in the network. Basal consumers (species that do not feed on others and serve as prey) are shown in panels (e-h). Panels (e, f) show in-degree, i.e., the number of predator and consumer species feeding on them, while panels (g, h) show eigenvector centrality, which reflects a species’ importance in the network by considering both its direct links and the centrality of its predators. Both trophic position and centrality reflect the mean over the species’ lifespan.

We detected phylogenetic signal in the Galápagos food webs, with a clear distinction between species roles. The strength of the signal varied with island characteristics associated with niche filling. For predator roles, the Mantel correlation between network and phylogenetic trait distances decreased with island age and increased with island area and elevation. For species in prey roles, the Mantel correlation also decreased with island age but showed the opposite direction for area and elevation, with correlations decreasing as area and elevation increased. Prey-role trends were based on 6 of the 13 islands where correlations were statistically significant (Figure 6a-c). The similarity between phylogenetic and network-based clusters (measured by NMI) increased with island age, in contrast to the Mantel test results. NMI also tended to increase with island area and elevation; however, the observed values were not significantly different from those generated under permutation (Figure 6d-f).

**Figure 6.**
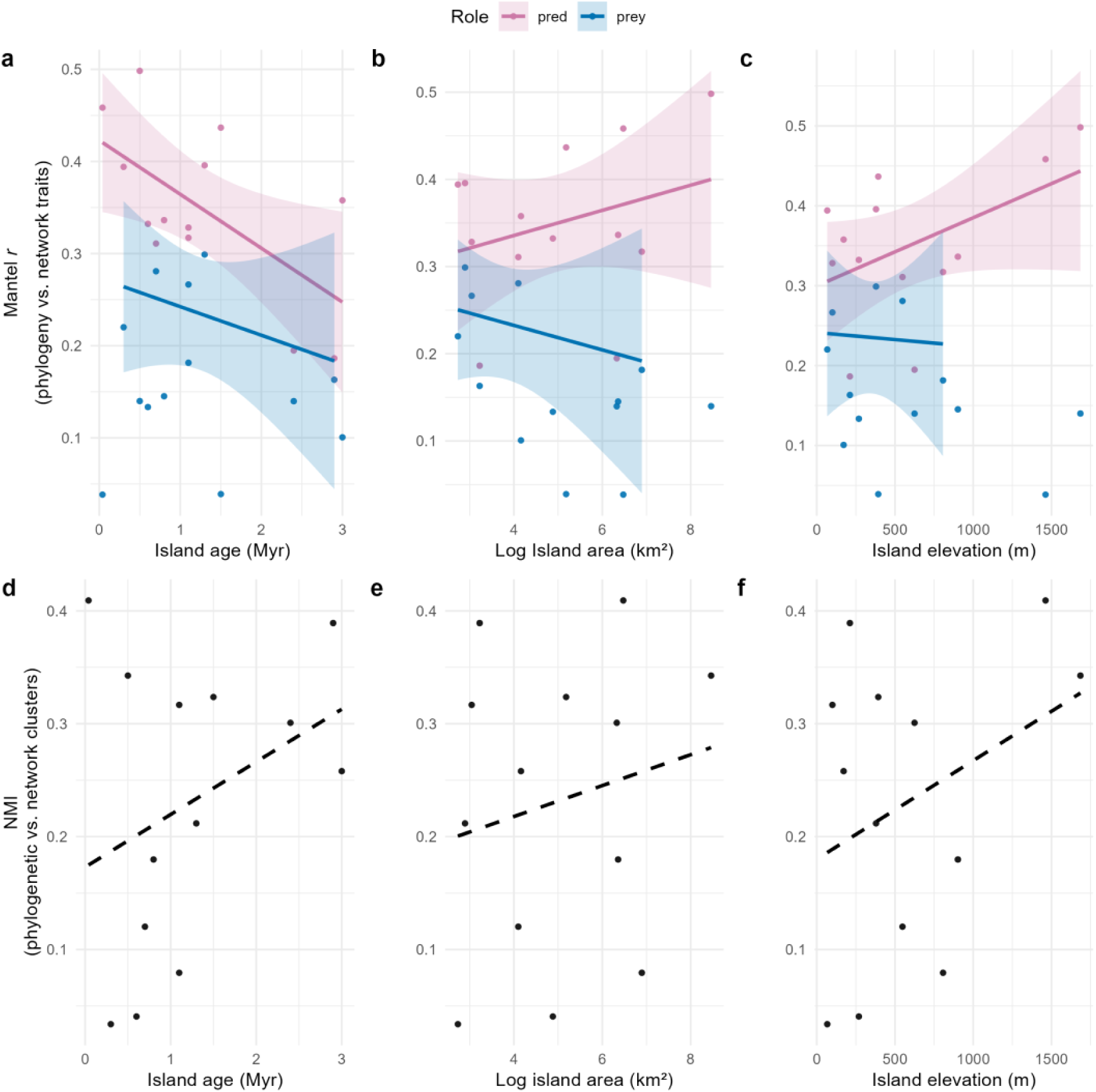
Phylogenetic signal in Galápagos vertebrate food webs. Phylogenetic signal based on two methods (panel rows) plotted against three island characteristics hypothesized to influence niche filling (panel columns). Island age reflects the stage of community assembly and the progressive reduction of unoccupied niche space over time. Island area and elevation represent habitat availability and topographic complexity, respectively, both promoting niche diversity. Each point represents one of the 13 islands studied. Filled dots indicate significant correlations based on randomization tests, while empty dots indicate non-significant results. Lines show fitted linear trends with 95% confidence bands (shaded areas). Solid lines consider only significant values in the fit.

The relationship between species’ phylogenetic distinctiveness (PD) and out-degree centrality (a proxy for diet breadth) varied across environmental gradients (Figure 7). It tended to weaken or become slightly negative with increasing island age. In contrast, positive trends were observed with island area and elevation. Most estimated regression coefficients were small in magnitude (|β| < 0.15), although some islands exhibited stronger relationships (range: -0.28 to 0.27).

**Figure 7.**
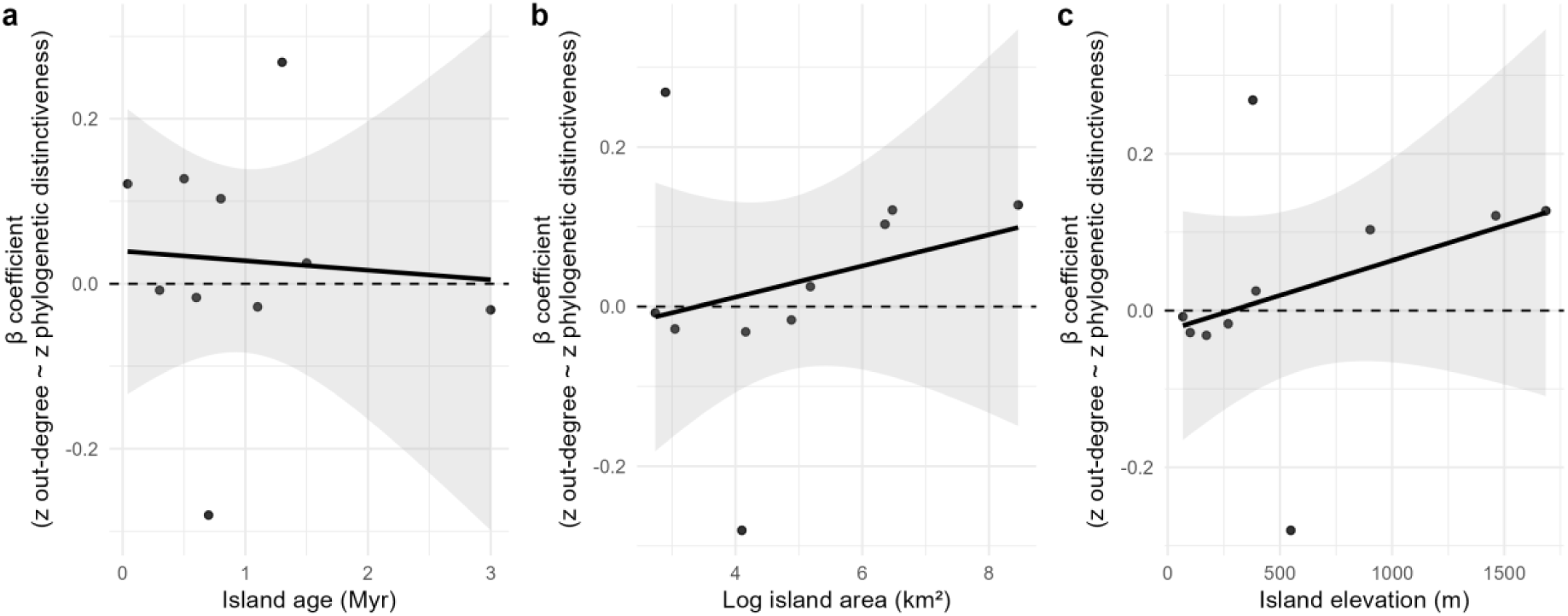
Relationship between phylogenetic distinctiveness and centrality in Galápagos vertebrate food webs. Plots show the island-level slope (β coefficient) from linear models relating species’ phylogenetic distinctiveness (PD) to out-degree centrality (a proxy for diet breadth) as a function of three island characteristics: (a) age, (b) area, and (c) elevation. Island age reflects the stage of community assembly and the progressive reduction of unoccupied niche space over time. Island area and elevation represent habitat availability and topographic complexity, respectively, both promoting niche diversity. Each point represents an island. Solid lines show fitted linear trends with 95% confidence bands (shaded areas). Positive slopes indicate that more phylogenetically distinct species tend to have broader diets (less dietary overlap with others), while negative slopes suggest that more phylogenetically distinct species have narrower or more specialized diets.

## Discussion

In this study, we explored the imprint of evolutionary history on ecological networks.. Using simulations that compare species’ positions in the network and phylogeny, we find evidence that phylogenetic signal declines as communities assemble and niche space becomes saturated. This erosion of signal appears to emerge from a broad set of feedbacks between ecological interactions, extinction, and the emergent structure of the food web. Species with high phylogenetic distinctiveness tend to be more specialized and occupy peripheral network positions. Diversification dynamics vary by trophic level and assembly stage: centrality may constrain diversification in intermediate consumers throughout, could limit top predators after niche saturation, and shows nonlinear effects in basal species depending on predation and connectivity. In Galápagos food webs, we find results that both align with and deviate from our predictions. Phylogenetic signal declines with island age, but responses to area and elevation differ between foraging and vulnerability traits, which may suggest that role-specific processes govern the persistence of phylogenetic structure.

The decline in correspondence between phylogeny and species’ network positions as communities diversify and niche space fills is consistent with classic theory. Early in assembly, new species resemble their ancestors and establish in unoccupied niches, preserving phylogenetic signal through niche conservatism or neutral evolution (Harvey & Pagel 1991, Price 1997, Blomberg & Garland 2002, Wiens & Graham 2005, Losos 2008). As niches saturate, competition promotes trait divergence among close relatives and convergence among distant ones, gradually eroding signal (Diamond 1975, Webb et al. 2002, Silvertown et al. 2006, Kraft et al. 2007, Cardillo 2011).

Our analysis suggests that the erosion of phylogenetic signal is not solely due to intensified competition, but may arise from broader network-mediated feedbacks involving extinction, diversification, and structural reorganization. Competition peaks early when niche overlap is high, then declines as the community matures. Extinction prunes redundant species, while trait divergence and specialization reduce overlap, potentially leading to a more modular, compartmentalized network. As modularity rises and connectance declines, species interact mainly within cohesive subgroups, diminishing broad-scale competition. Rather than sustaining competition, niche saturation appears to trigger a self-organizing process that limits its influence through network reconfiguration and ecological differentiation.

This structural reorganization may further weaken the link between phylogeny and ecological roles. Close relatives diverge functionally as they occupy different modules or roles, while distantly related species converge on similar roles. Most critically, extinction is shaped by interacting trophic pressures—competition, predation, and resource limitation—that act across clades in complex, non-additive ways. As a result, extinctions are scattered across the phylogeny, further diluting phylogenetic structure in network positions.

Nonetheless, our model is a simplified system explored within a limited parameter space, and should be seen as a heuristic tool and first approximation to uncover how feedbacks between diversification, interactions, and extinction influence phylogeny-network correspondence. While our simulations include randomness, they are largely driven by biotic interactions. In reality, stochastic events and abiotic conditions also shape assembly. For example, habitat filtering may promote phylogenetic clustering by favoring similar lineages (Webb et al. 2002), and random perturbations could remove clades or open new niche space through extinction (Cavender-Bares et al. 2009). Furthermore, our model assumes a closed community, while real ecosystems are often open to invasions. Depending on their traits and timing, invaders can either reinforce or disrupt phylogenetic structure (Proches et al. 2008). We also omit spatial dynamics and variation in the regional species pool, which are increasingly recognized as key drivers of phylogenetic signals in ecological networks (Maitner et al. 2022, Calcagno et al. 2023).

Finally, an important open challenge is to better link microevolutionary mechanisms of speciation with macroevolutionary patterns of community assembly (Harmon et al. 2019). This includes integrating demographic processes, trait evolution, and some reproductive isolation mechanisms. One promising direction is to couple metacommunity models that already incorporate speciation via genetic incompatibilities—such as the Bateson-Dobzhansky-Muller framework used by Desjardins-Proulx et al. (2012)—with the evolution of ecological traits that mediate species interactions. This would allow ecological divergence to influence reproductive isolation, for instance, through pleiotropic links between ecological function and genetic incompatibility (Rundle & Nosil 2005, Gavrilets 2014, Kulmuni & Westram 2017).

Despite its simplicity, our model captures processes that likely shape community assembly across systems. We predict that phylogeny-network correspondence declines as niche space fills, with stronger phylogenetic signal in young, species-poor communities undergoing early assembly or ecological opportunity, and weaker signal in mature, species-rich systems where ecological roles have diverged. This prediction mirrors observed contrasts between island and continental systems: island biotas, shaped by limited colonization and in situ speciation, often show phylogenetic clustering (Losos & Schluter 2000, Si et al. 2022), while continental systems—with greater biotic resistance and less adaptive trait evolution—tend to show phylogenetic overdispersion (Proches et al. 2008).

These dynamics may also extend to island-like systems, such as fragmented landscapes, where niche availability similarly structures communities. For instance, phylogenetic signal in pollination networks has been shown to increase with fragment area (Aizen et al. 2016). We also expect scale-dependent competitive exclusion may reduce phylogenetic signal locally, while shared environmental filters promote co-occurrence of relatives regionally (Proches et al. 2008, Hutchinson et al. 2017). Finally, in transient or disturbance-prone environments, rapid extinction–recolonization dynamics may temporarily elevate phylogenetic signal, as clade-specific traits enable swift colonization of vacant niches— paralleling our simulation results.

In Galápagos vertebrate food webs, we find that phylogenetic signal in species’ foraging and vulnerability roles declines with island age, broadly consistent with expectations from niche filling and network reorganization. However, for the island area and elevation, we observe contrasting trends between predator (increases) and prey (declines) roles, which may reflect role-specific selective pressures. On larger, higher islands, predators may retain conserved foraging traits due to increased niche availability and reduced competition (Losos & Schluter 2000) while prey likely experience stronger enemy pressure, driving divergence in vulnerability traits like defenses or microhabitat use (Johnson & Belk 2020). Greater lability in these vulnerability traits may further accelerate this divergence. While multiple processes may generate similar signals (Cardillo et al. 2008, Gerhold et al. 2015), our results highlight that differences in the ecological roles of species may mediate the strength and direction of phylogenetic signal under varying environmental and community contexts, asymmetries also described in other antagonistic networks (Bersier & Kehrli 2008, Krasnov et al. 2012, Rohr & Bascompte 2014).

Although not significant, the clustering analysis revealed discrepancies with the Mantel test for island age, underscoring the need for standardized methods to quantify phylogeny-network correspondence. Most existing approaches rely on pairwise correlations between phylogenetic distances and interaction similarity (e.g., Bersier & Kehrli 2008, Elias et al. 2013, Perez-lamarque et al. 2022), or GLS models accounting for shared ancestry (Ives & Godfray 2006), but they overlook the multivariate structure of networks and phylogenies. Here, we extend the approach of Dalla Riva & Stouffer (2015), embedding species into multivariate trait spaces based on their topological roles and comparing these with phylogenetic embeddings. This treats phylogeny-network alignment as a higher-dimensional problem and has been applied in other systems (Joffard et al. 2019, Massol et al. 2021). While some studies explore phylogenetic signal in modularity or node-level traits (e.g., Poulin et al. 2013), general multivariate methods remain underdeveloped.

As communities assemble, phylogenetically distinct species tend to be more specialized and occupy peripheral network positions, possibly due to difficulties competing with an already diversified network core. This may suggest that early-diversifying clades form a central, stable core that is harder to invade, while later-evolving species occupy remaining, more marginal niches—leading to peripheral positions in the network. While it has been proposed that evolutionarily distinct species may play unique or complementary roles (Davies et al. 2016), recent analyses show they often have fewer interactions (Kruger & Davies 2025), consistent with our results. In Galápagos vertebrate food webs, the relationship between phylogenetic distinctiveness and diet breadth was weakly positive overall—possibly due to introduced generalists like rats and cats—but turned negative under island characteristics associated with niche filling, which may support the idea that newer species face increasing barriers to centrality as networks mature. Further research is needed to clarify these mechanisms. In addition, because our network relies on a metaweb approach—assuming interactions whenever species co-occur based on compiled literature—it may miss local specialization (Blanchet et al. 2020). A necessary step is to revise the network using local interaction data.

We found that the relationship between species’ network positions and diversification varies by trophic level and the stage of community assembly. For both top predators and intermediate consumers, diversification rates declined with increasing generalism (i.e., in-degree) before niche saturation. At this stage, network connectance is larger, and a possible explanation for this relationship is that connectivity exposes consumers to greater ecological constraints: in intermediate consumers, this likely reflects a combined effect of increased resource overlap (competition) and exposure to top-down control from predators, which may suppress diversification. Top predators, not being preyed upon, are likely more affected by competition alone. Notably, betweenness centrality also showed a consistent negative relationship with diversification in intermediate consumers during this phase, suggesting that occupying bridging positions across the network may impose costs through unstable or conflicting ecological pressures. After niche saturation, the network undergoes a structural reorganization, becoming more modular. In this context, the relationship between generalism and diversification becomes hump-shaped for both top predators and intermediate consumers. This shift suggests that extreme strategies—either high specialization or broad generalism—are disfavored. Specialists may face instability due to narrow diet breadth and the risk of losing key resources, while highly generalist consumers may encounter intensified competition within modules. Importantly, betweenness centrality no longer predicts diversification in this modular phase. This may be because bridging across modules becomes less ecologically relevant as compartments stabilize interaction flows and limit the structural consequences of cross-module positions. Species no longer act as key connectors between distant parts of the network, reducing the ecological costs or opportunities associated with being a structural bridge.

For basal consumers, diversification declined with increasing out-degree (i.e., number of consumers), indicating that being heavily preyed upon may constrain diversification opportunities, possibly due to higher extinction risks or reduced ecological opportunity. Interestingly, eigenvector centrality showed a hump-shaped relationship, with diversification peaking at intermediate to high centrality, suggesting that basal species connected to influential consumers may benefit from indirect effects—such as diffuse predation or release from competition—that promote divergence and speciation. The integration of other interaction types, such as facilitation, could substantially alter community assembly dynamics and should be included in future models (Fontaine et al. 2011, Pedraza et al. 2024).

Testing these predictions in empirical systems requires linking local interaction networks with the biogeographic and evolutionary context of the clades involved. Such integrative data are increasingly available for well-studied groups, enabling deeper exploration of how macroevolution shapes species’ network roles. For instance, our prediction that intermediate consumers have lower speciation rates in saturated communities—especially generalists— partly matches global analysis of avian diets, which show that omnivorous birds experience both higher extinction and lower speciation rates (Burin et al. 2016), and that clades with higher diversification or lower extinction contributed more central species to seed-dispersal networks (Burin et al. 2021). A promising avenue to explore these questions is through sampling interaction networks across island radiations, which offer rich phylogenetic data within a well-defined biogeographic framework (Emerson 2002).

Our findings advance a mechanistic understanding of how evolutionary history becomes uncoupled from network structure during community assembly. By linking diversification, ecological interactions, and network reorganization, we suggest that phylogenetic signal is transient and context-dependent. Trophic feedbacks and niche availability play key roles in shaping the macroevolutionary imprint of food webs. Advancing this framework will require models that incorporate spatial dynamics and diverse interactions, along with empirical studies and improved methods for comparing phylogenies and network structures.

## Acknowledgements

We thank the Quebec Centre for Biodiversity Science (QCBS) and the Computational Biodiversity Science and Services (BIOS²) training program for supporting AFC’s research stays at the Institut Pasteur de Lille and the University of São Paulo. Financial support was provided by the NSERC - CREATE Training program in computational biodiversity science and the NSERC Discovery Grant to DG.

