## Supplement 1 for "Phylogenetic signal dynamics during niche filling in food webs"

**Table S1. Model parameters.**

| Parameter | Description | Value |
|---|---|---|
| $n_i$ | Niche position of species $i$ [0,1]; initialized $U(0{,}1, 0.3)$ | |
| $r_i$ | Niche breadth (half-range) of species $i$ (>0); initialized $U(0{,}05, 0.15)$ | |
| $c_j$ | Foraging optimum of consumer j | $c_j = n_j/2$ |
| L | Trophic interaction matrix at a timestep | Variable |
| S | Number of extant non-basal species at the timestep | Variable |
| $S_{basal}$ | Basal pool consisting on 5 producers; niche positions $U(0, 0.3)$ | |
| $d_{ij}$ | Trait distance for interaction: $d_{ij} = \lvert n_i - c_j \rvert$ | Variable |
| $D_{out}$ | Number of predators of a focal species | Variable |
| $I_{out}$ | Number of predators consuming a herbivore | Variable |
| $S^{pred}$ | Predator similarity matrix | Variable |
| $\phi_i$ | Avg dietary similarity of predator $i$ | Variable |
| $\sigma$ | Controls decay of interaction probability with niche distance | 0.04 |
| $p_{up}$ | Penalty factor for upward interactions | 0.1 |

| | | |
|---|---|---|
| | (prey niche > predator niche) | |
| $u_{max}$ | Maximum speciation rate at low diversity | 0.3 |
| $\delta$ | Strength of diversity dependence on speciation | 0.5 |
| $I_{max}$ | Species richness threshold for niche saturation | 60 |
| $m_{scale}$ | Scaling factor for trait displacement during speciation | 0.7 |
| $u_0$ | Establishment propensity at high predation | 0.2 |
| $u_1$ | Establishment propensity at zero predation | 1.2 |
| $a_u$ | Rate of decline in establishment with predator numbe | 0.3 |
| $e_0$ | Extinction propensity at high predation | 0.1 |
| $e_1$ | Extinction propensity at zero predation | 0.3 |
| $a_e$ | Rate of increase in extinction with predator number | 0.6 |
| $\theta_n$ | Minimum niche difference required to avoid competitive exclusion of the ancestor | 0.05 |
| $\kappa$ | Sensitivity to competitive overlap among predators | 0.2 |
| $B_{ext}$ | Weight of top-down vs competition extinction | 0.5 |

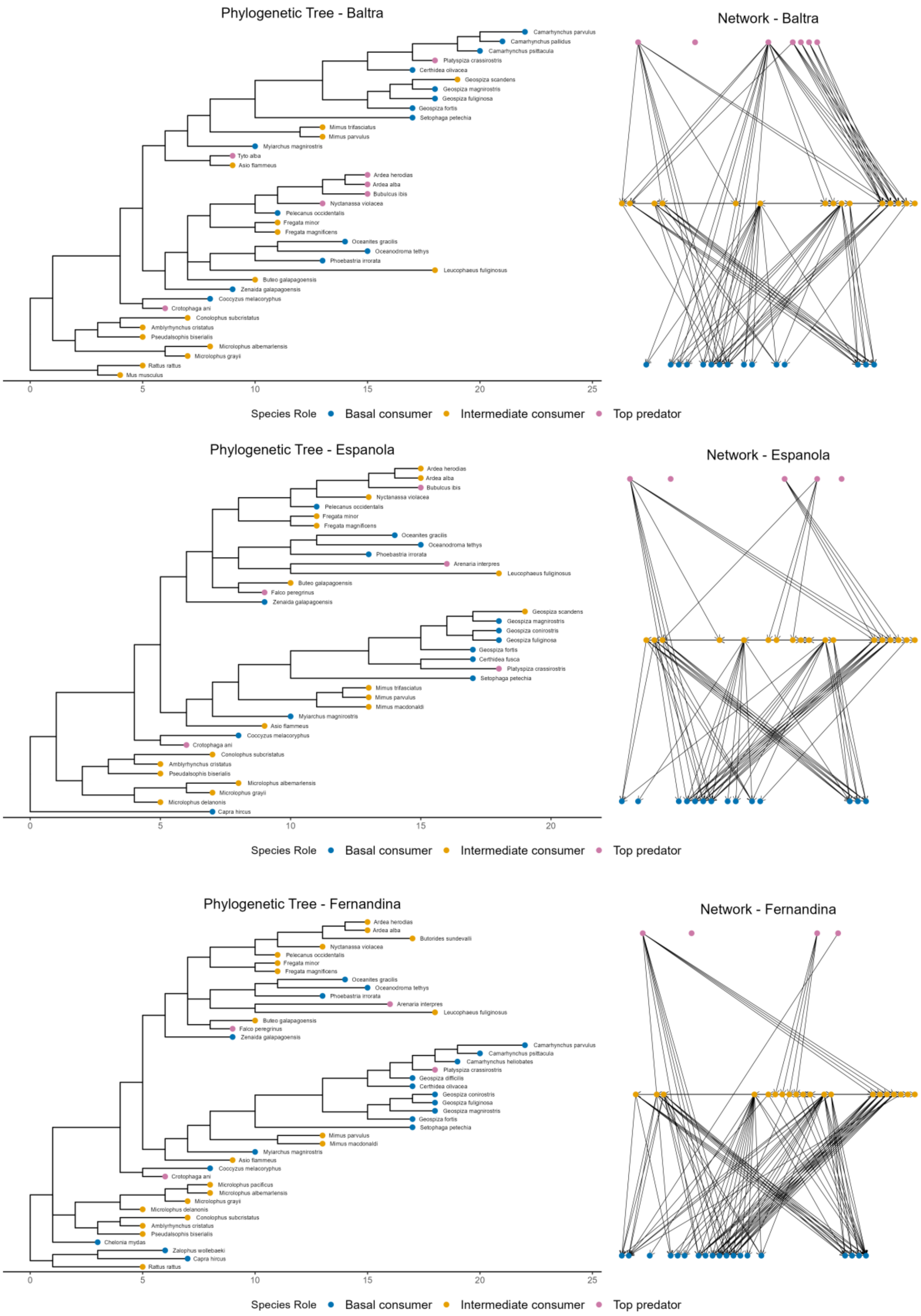

Phylogenetic Tree - Baltra
Camarhynchus parvulus
Camarhynchus pallidus
Camarhynchus psittacula
Platyspiza crassirostris
Certhidea olivacea
Geospiza scandens
Geospiza magnirostris
Geospiza fuliginosa
Geospiza fortis
Setophaga petechia
Mimus trifasciatus
Mimus parvulus
Myiarchus magnirostris
Tyto alba
Asio flammeus
Ardea herodias
Ardea alba
Bubulcus ibis
Nyctanassa violacea
Pelecanus occidentalis
Fregata minor
Fregata magnificens
Oceanites gracilis
Oceanodroma tethys
Phoebastria irrorata
Leucophaeus fuliginosus
Buteo galapagoensis
Zenaida galapagoensis
Coccyzus melacoryphus
Crotophaga ani
Conolophus subcristatus
Amblyrhynchus cristatus
Pseudalsophis biserialis
Microlophus albemarlensis
Microlophus grayii
Rattus rattus
Mus musculus
0
5
10
15
20
25
Network - Baltra
Species Role
Basal consumer
Intermediate consumer
Top predator
Phylogenetic Tree - Espanola
Ardea herodias
Ardea alba
Bubulcus ibis
Nyctanassa violacea
Pelecanus occidentalis
Fregata minor
Fregata magnificens
Oceanites gracilis
Oceanodroma tethys
Phoebastria irrorata
Arenaria interpres
Leucophaeus fuliginosus
Buteo galapagoensis
Falco peregrinus
Zenaida galapagoensis
Geospiza scandens
Geospiza magnirostris
Geospiza conirostris
Geospiza fuliginosa
Geospiza fortis
Certhidea fusca
Platyspiza crassirostris
Setophaga petechia
Mimus trifasciatus
Mimus parvulus
Mimus macdonaldi
Myiarchus magnirostris
Asio flammeus
Coccyzus melacoryphus
Crotophaga ani
Conolophus subcristatus
Amblyrhynchus cristatus
Pseudalsophis biserialis
Microlophus albemarlensis
Microlophus grayii
Microlophus delanonis
Capra hircus
0
5
10
15
20
Network - Espanola
Species Role
Basal consumer
Intermediate consumer
Top predator
Phylogenetic Tree - Fernandina
Ardea herodias
Ardea alba
Butorides sundevalli
Nyctanassa violacea
Pelecanus occidentalis
Fregata minor
Fregata magnificens
Oceanites gracilis
Oceanodroma tethys
Phoebastria irrorata
Arenaria interpres
Leucophaeus fuliginosus
Buteo galapagoensis
Falco peregrinus
Zenaida galapagoensis
Camarhynchus parvulus
Camarhynchus psittacula
Camarhynchus heliobates
Platyspiza crassirostris
Geospiza difficilis
Certhidea olivacea
Geospiza conirostris
Geospiza fuliginosa
Geospiza magnirostris
Geospiza fortis
Setophaga petechia
Mimus parvulus
Mimus macdonaldi
Myiarchus magnirostris
Asio flammeus
Coccyzus melacoryphus
Crotophaga ani
Microlophus pacificus
Microlophus albemarlensis
Microlophus grayii
Microlophus delanonis
Conolophus subcristatus
Amblyrhynchus cristatus
Pseudalsophis biserialis
Chelonia mydas
Zalophus wollebaeki
Capra hircus
Rattus rattus
0
5
10
15
20
25
Network - Fernandina
Species Role
Basal consumer
Intermediate consumer
Top predator

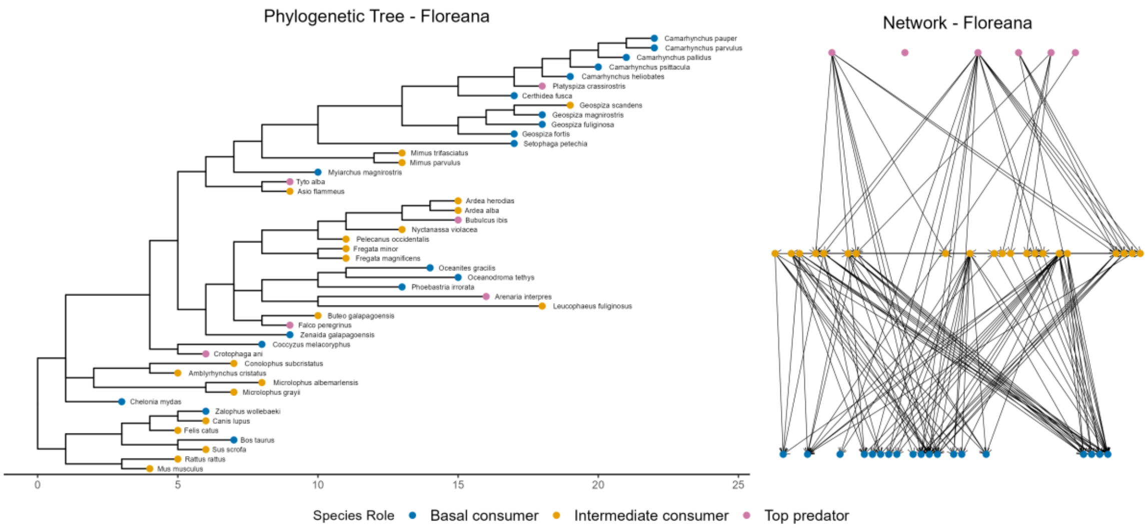
Phylogenetic Tree - Floreana
Network - Floreana
Camarhynchus pauper
Camarhynchus parvulus
Camarhynchus pallidus
Camarhynchus psittacula
Camarhynchus heliobates
Platyspiza crassirostris
Certhidea fusca
Geospiza scandens
Geospiza magnirostris
Geospiza fuliginosa
Geospiza fortis
Setophaga petechia
Mimus trifasciatus
Mimus parvulus
Myiarchus magnirostris
Tyto alba
Asio flammeus
Ardea herodias
Ardea alba
Bubulcus ibis
Nyctanassa violacea
Pelecanus occidentalis
Fregata minor
Fregata magnificens
Oceanites gracilis
Oceanodroma tethys
Phoebastria irrorata
Arenaria interpres
Leucophaeus fuliginosus
Buteo galapagoensis
Falco peregrinus
Zenaida galapagoensis
Coccyzus melacoryphus
Crotophaga ani
Conolophus subcristatus
Amblyrhynchus cristatus
Microlophus albemarlensis
Microlophus grayii
Chelonia mydas
Zalophus wollebaeki
Canis lupus
Felis catus
Bos taurus
Sus scrofa
Rattus rattus
Mus musculus
0
5
10
15
20
25
Species Role
Basal consumer
Intermediate consumer
Top predator

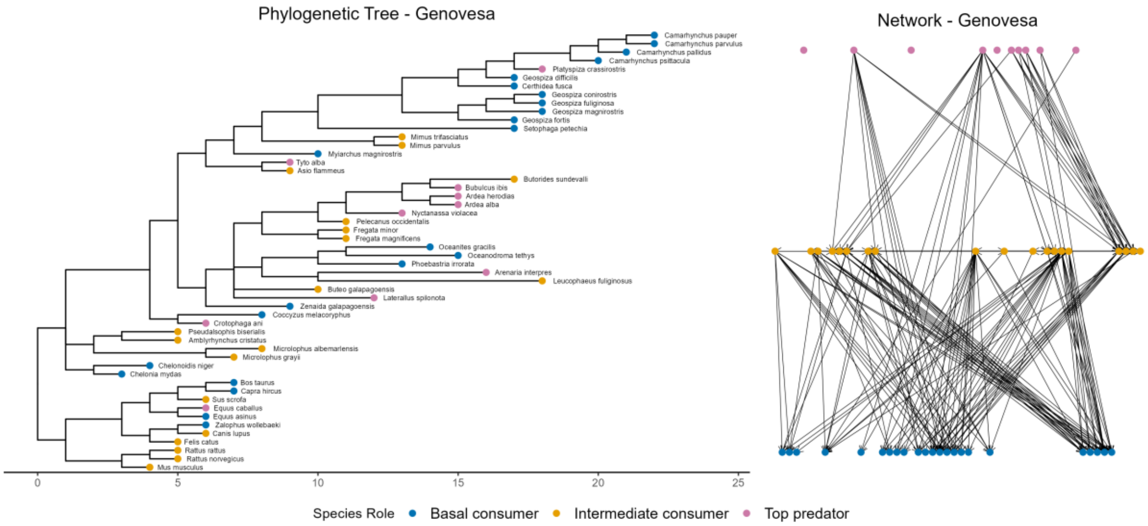
Phylogenetic Tree - Genovesa
Network - Genovesa
Camarhynchus pauper
Camarhynchus parvulus
Camarhynchus pallidus
Camarhynchus psittacula
Platyspiza crassirostris
Geospiza difficilis
Certhidea fusca
Geospiza conirostris
Geospiza fuliginosa
Geospiza magnirostris
Geospiza fortis
Setophaga petechia
Mimus trifasciatus
Mimus parvulus
Myiarchus magnirostris
Tyto alba
Asio flammeus
Butorides sundevalli
Bubulcus ibis
Ardea herodias
Ardea alba
Nyctanassa violacea
Pelecanus occidentalis
Fregata minor
Fregata magnificens
Oceanites gracilis
Oceanodroma tethys
Phoebastria irrorata
Arenaria interpres
Leucophaeus fuliginosus
Buteo galapagoensis
Laterallus spilonota
Zenaida galapagoensis
Coccyzus melacoryphus
Crotophaga ani
Pseudalsophis biserialis
Amblyrhynchus cristatus
Microlophus albemarlensis
Microlophus grayii
Chelonoidis niger
Chelonia mydas
Bos taurus
Capra hircus
Sus scrofa
Equus caballus
Equus asinus
Zalophus wollebaeki
Canis lupus
Felis catus
Rattus rattus
Rattus norvegicus
Mus musculus
0
5
10
15
20
25
Species Role
Basal consumer
Intermediate consumer
Top predator

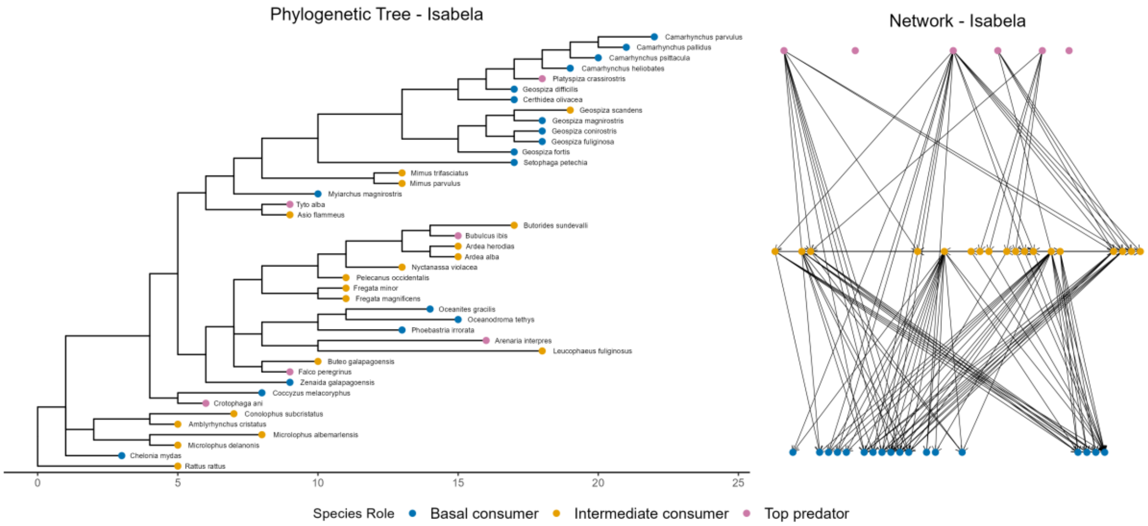
Phylogenetic Tree - Isabela
Network - Isabela
Camarhynchus parvulus
Camarhynchus pallidus
Camarhynchus psittacula
Camarhynchus heliobates
Platyspiza crassirostris
Geospiza difficilis
Certhidea olivacea
Geospiza scandens
Geospiza magnirostris
Geospiza conirostris
Geospiza fuliginosa
Geospiza fortis
Setophaga petechia
Mimus trifasciatus
Mimus parvulus
Myiarchus magnirostris
Tyto alba
Asio flammeus
Butorides sundevalli
Bubulcus ibis
Ardea herodias
Ardea alba
Nyctanassa violacea
Pelecanus occidentalis
Fregata minor
Fregata magnificens
Oceanites gracilis
Oceanodroma tethys
Phoebastria irrorata
Arenaria interpres
Leucophaeus fuliginosus
Buteo galapagoensis
Falco peregrinus
Zenaida galapagoensis
Coccyzus melacoryphus
Crotophaga ani
Conolophus subcristatus
Amblyrhynchus cristatus
Microlophus albemarlensis
Microlophus delanonis
Chelonia mydas
Rattus rattus
0
5
10
15
20
25
Species Role
Basal consumer
Intermediate consumer
Top predator

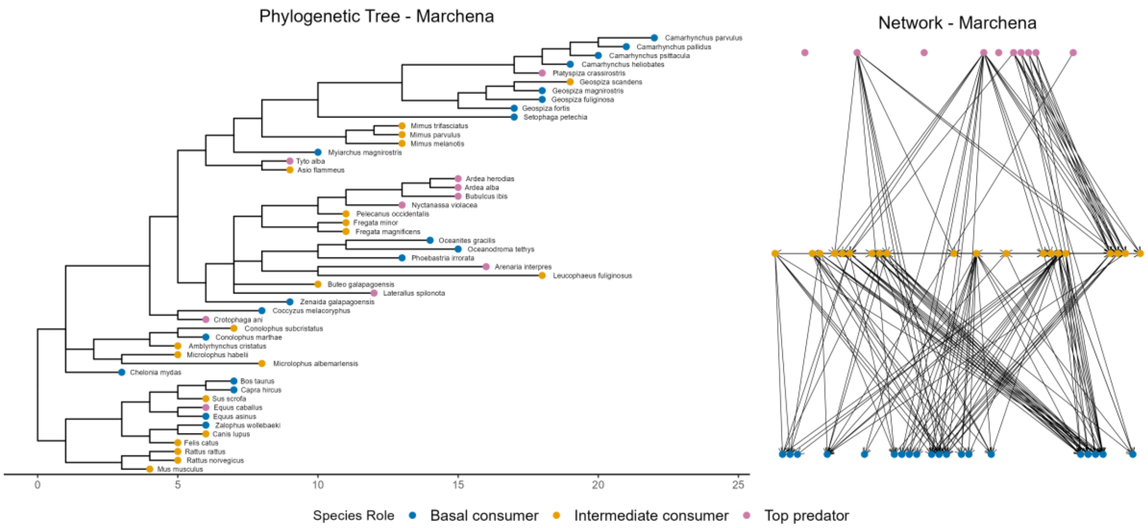
Phylogenetic Tree - Marchena
Camarhynchus parvulus
Camarhynchus pallidus
Camarhynchus psittacula
Camarhynchus heliobates
Platyspiza crassirostris
Geospiza scandens
Geospiza magnirostris
Geospiza fuliginosa
Geospiza fortis
Setophaga petechia
Mimus trifasciatus
Mimus parvulus
Mimus melanotis
Myiarchus magnirostris
Tyto alba
Asio flammeus
Ardea herodias
Ardea alba
Bubulcus ibis
Nyctanassa violacea
Pelecanus occidentalis
Fregata minor
Fregata magnificens
Oceanites gracilis
Oceanodroma tethys
Phoebastria irrorata
Arenaria interpres
Leucophaeus fuliginosus
Buteo galapagoensis
Laterallus spilonota
Zenaida galapagoensis
Coccyzus melacoryphus
Crotophaga ani
Conolophus subcristatus
Conolophus marthae
Amblyrhynchus cristatus
Microlophus habelii
Microlophus albemarlensis
Chelonia mydas
Bos taurus
Capra hircus
Sus scrofa
Equus caballus
Equus asinus
Zalophus wollebaeki
Canis lupus
Felis catus
Rattus rattus
Rattus norvegicus
Mus musculus
0
5
10
15
20
25
Network - Marchena
Species Role
Basal consumer
Intermediate consumer
Top predator

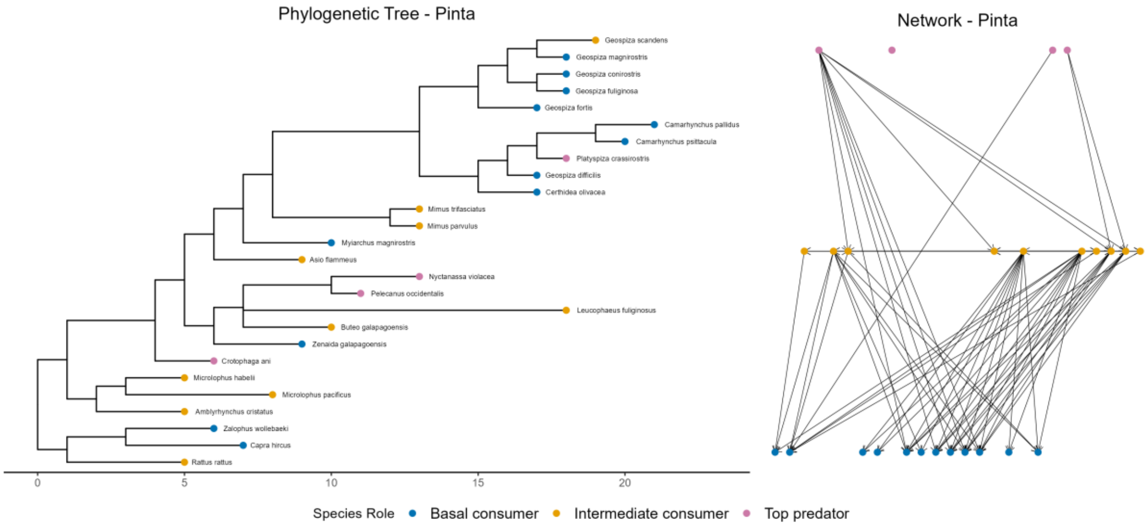
Phylogenetic Tree - Pinta
Geospiza scandens
Geospiza magnirostris
Geospiza conirostris
Geospiza fuliginosa
Geospiza fortis
Camarhynchus pallidus
Camarhynchus psittacula
Platyspiza crassirostris
Geospiza difficilis
Certhidea olivacea
Mimus trifasciatus
Mimus parvulus
Myiarchus magnirostris
Asio flammeus
Nyctanassa violacea
Pelecanus occidentalis
Leucophaeus fuliginosus
Buteo galapagoensis
Zenaida galapagoensis
Crotophaga ani
Microlophus habelii
Microlophus pacificus
Amblyrhynchus cristatus
Zalophus wollebaeki
Capra hircus
Rattus rattus
0
5
10
15
20
Network - Pinta
Species Role
Basal consumer
Intermediate consumer
Top predator

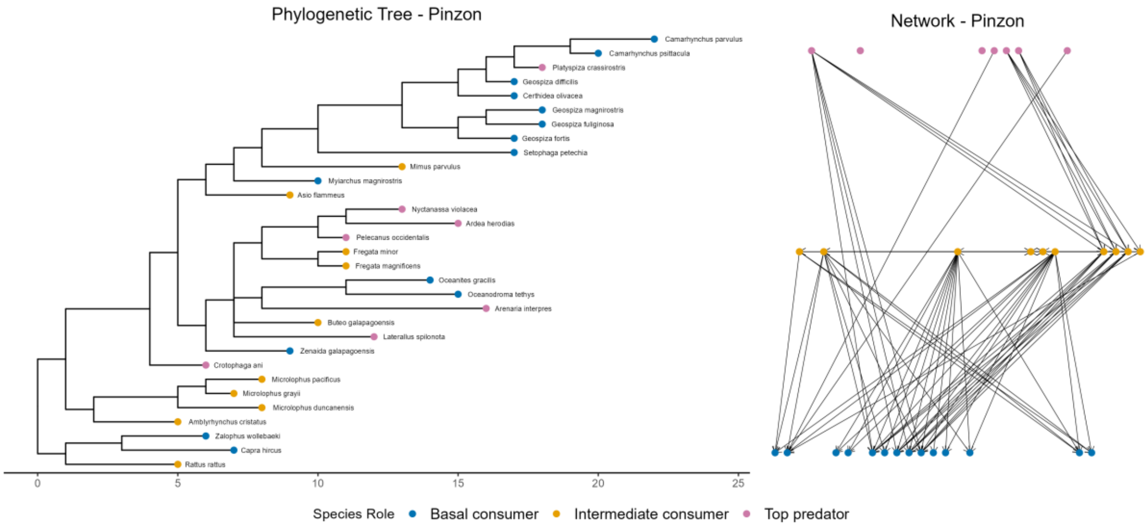
Phylogenetic Tree - Pinzon
Camarhynchus parvulus
Camarhynchus psittacula
Platyspiza crassirostris
Geospiza difficilis
Certhidea olivacea
Geospiza magnirostris
Geospiza fuliginosa
Geospiza fortis
Setophaga petechia
Mimus parvulus
Myiarchus magnirostris
Asio flammeus
Nyctanassa violacea
Ardea herodias
Pelecanus occidentalis
Fregata minor
Fregata magnificens
Oceanites gracilis
Oceanodroma tethys
Arenaria interpres
Buteo galapagoensis
Laterallus spilonota
Zenaida galapagoensis
Crotophaga ani
Microlophus pacificus
Microlophus grayii
Microlophus duncanensis
Amblyrhynchus cristatus
Zalophus wollebaeki
Capra hircus
Rattus rattus
0
5
10
15
20
25
Network - Pinzon
Species Role
Basal consumer
Intermediate consumer
Top predator

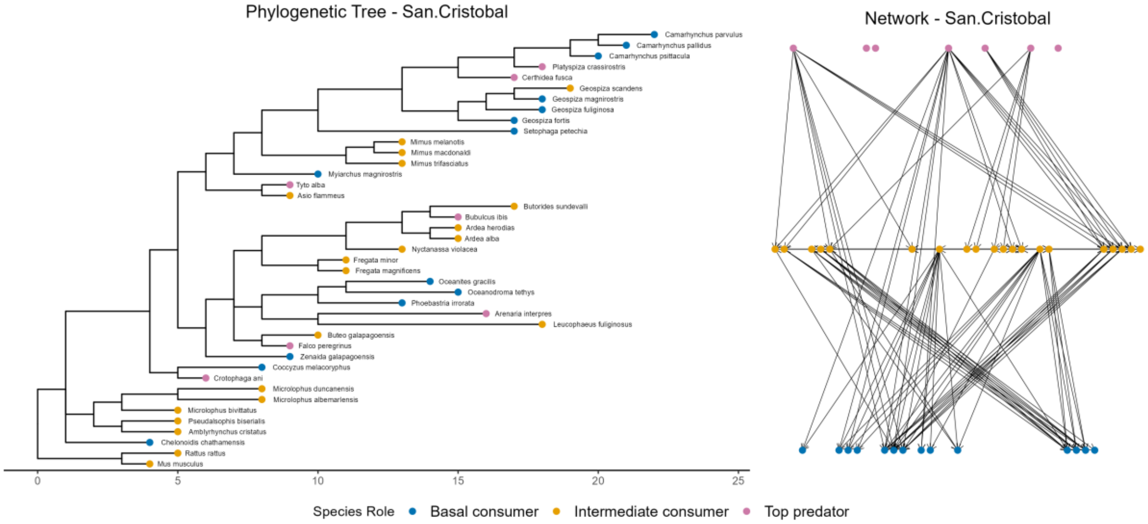

Phylogenetic Tree - San.Cristobal
Network - San.Cristobal
0 5 10 15 20 25
Species Role
Basal consumer
Intermediate consumer
Top predator


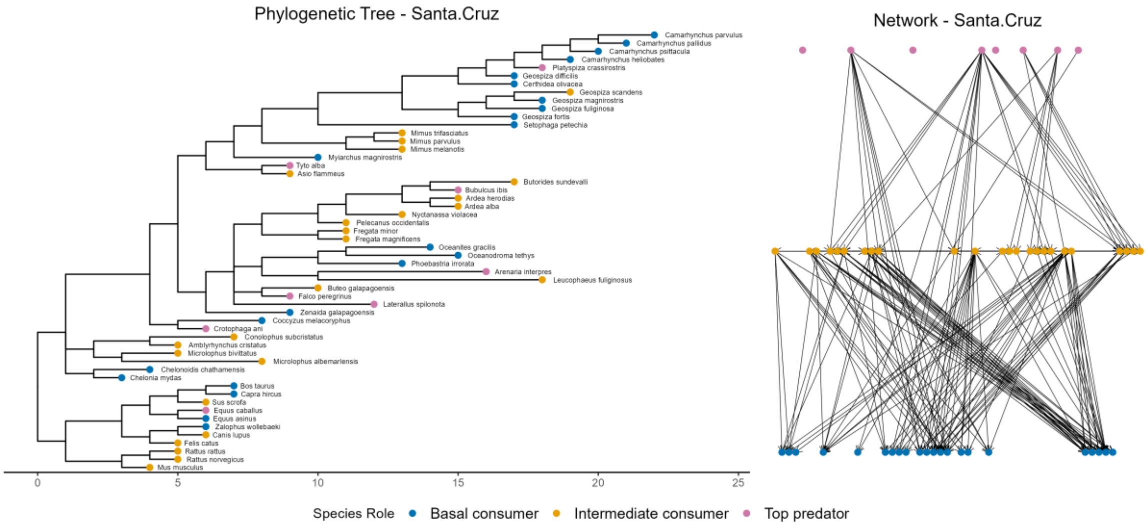

Phylogenetic Tree - Santa.Cruz
Network - Santa.Cruz
0 5 10 15 20 25
Species Role
Basal consumer
Intermediate consumer
Top predator


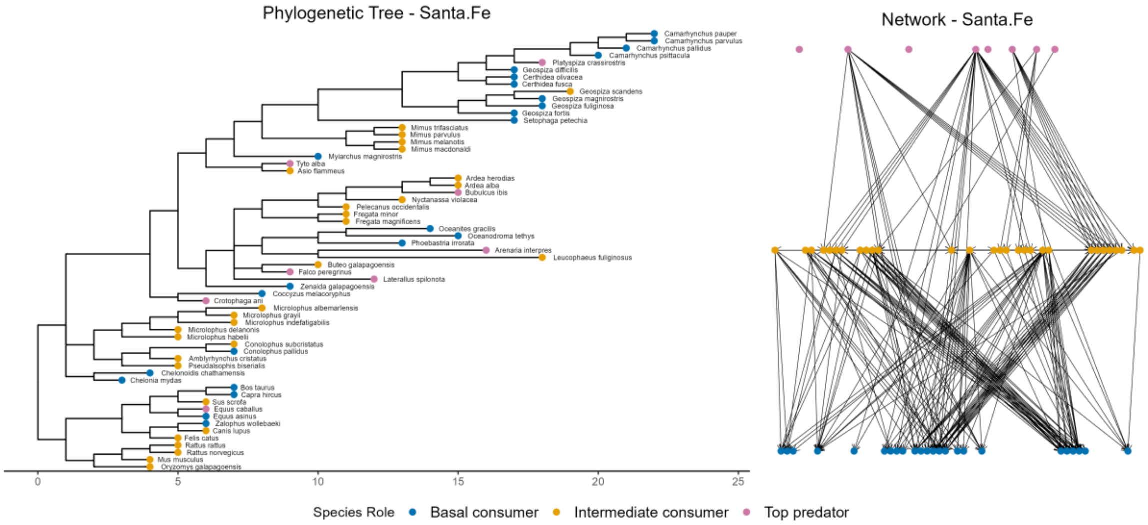

Phylogenetic Tree - Santa.Fe
Network - Santa.Fe
0 5 10 15 20 25
Species Role
Basal consumer
Intermediate consumer
Top predator

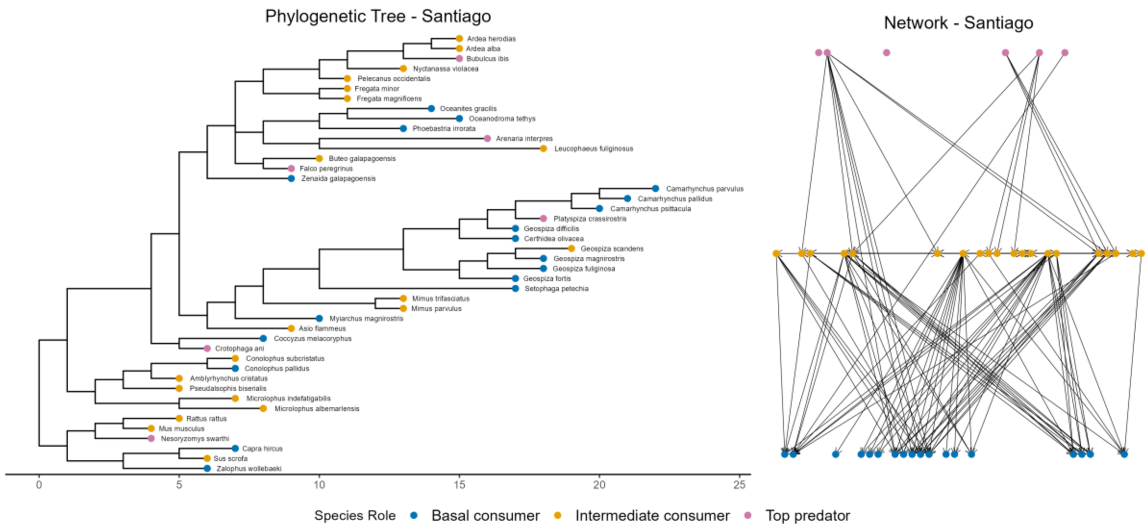


**Figure S1. Phylogenetic and network data analyzed per island.**
